# Rapid reconstruction of SARS-CoV-2 using a synthetic genomics platform

**DOI:** 10.1101/2020.02.21.959817

**Authors:** Tran Thi Nhu Thao, Fabien Labroussaa, Nadine Ebert, Philip V’kovski, Hanspeter Stalder, Jasmine Portmann, Jenna Kelly, Silvio Steiner, Melle Holwerda, Annika Kratzel, Mitra Gultom, Laura Laloli, Linda Hüsser, Manon Wider, Stephanie Pfaender, Dagny Hirt, Valentina Cippà, Silvia Crespo-Pomar, Simon Schröder, Doreen Muth, Daniela Niemeyer, Marcel A. Müller, Christian Drosten, Ronald Dijkman, Joerg Jores, Volker Thiel

## Abstract

Reverse genetics has been an indispensable tool revolutionising our insights into viral pathogenesis and vaccine development. Large RNA virus genomes, such as from Coronaviruses, are cumbersome to clone and to manipulate in *E. coli* hosts due to size and occasional instability^1-3^. Therefore, an alternative rapid and robust reverse genetics platform for RNA viruses would benefit the research community. Here we show the full functionality of a yeast-based synthetic genomics platform for the genetic reconstruction of diverse RNA viruses, including members of the *Coronaviridae, Flaviviridae* and *Paramyxoviridae* families. Viral subgenomic fragments were generated using viral isolates, cloned viral DNA, clinical samples, or synthetic DNA, and reassembled in one step in *Saccharomyces cerevisiae* using transformation associated recombination (TAR) cloning to maintain the genome as a yeast artificial chromosome (YAC). T7-RNA polymerase has been used to generate infectious RNA, which was then used to rescue viable virus. Based on this platform we have been able to engineer and resurrect chemically-synthetized clones of the recent epidemic SARS-CoV-2^4^ in only a week after receipt of the synthetic DNA fragments. The technical advance we describe here allows to rapidly responding to emerging viruses as it enables the generation and functional characterization of evolving RNA virus variants - in real-time - during an outbreak.

## Main Text

Population growth, increased travel and climate change foster epidemic and pandemic threats by (re)emerging RNA viruses. Although black swan events cannot be predicted, our medical sector should be ready to use the latest advancements in science to combat emerging viruses. Within the past decade we have seen outbreaks of Middle-East Respiratory Syndrome coronavirus (MERS-CoV)^5^, ZIKA virus (ZIKV)^6^, Ebola virus^7^, and at the end of 2019, SARS-CoV-2 that was first detected in Wuhan, Hubei province, China^4^, but meanwhile spread throughout China and beyond. Until now SARS-CoV-2 isolates from China have not been shared with health authorities and the scientific community, although they are urgently needed to build up serological diagnostics, to develop and assess antivirals and vaccines, and to establish appropriate *in vivo* models. Generation of SARS-CoV-2 from chemically synthesized DNA could bypass the limited availability of virus isolates and would furthermore allow genetic modifications and functional characterization of individual genes. However, although the workhorse *E. coli* did prove very useful for cloning many viral genomes, it has drawbacks for a number of emerging RNA viruses, including coronaviruses, in terms of assembling and stably maintaining full-length molecular clones.

Synthetic genomics is a field that was fuelled by the efforts to create a bacterial cell that is controlled by a synthetic genome ^8^. Genome-wide reassembly of the ∼1.1Mb *Mycoplasma* genome was first attempted using *E. coli* as an intermediate host ^8^, but the maintenance of these 100-kbp DNA fragments appeared to be very difficult in this host. Therefore, the yeast *Saccharomyces cerevisiae* was chosen to clone, assemble and even mutagenize entire *Mycoplasma* genomes ^9,10^. This ultimately led to the construction of the Mycoplasma JCVI Syn3.0, a cell carrying the minimal set of genes allowing its autonomous replication in auxotrophic medium ^11^. The rationale for using a yeast cloning system is the ability of yeast to recombine overlapping DNA fragments *in cellulo*, which led to the development of a technique called “transformation-associated recombination” (TAR) cloning^12^.

More recently, TAR cloning was successfully applied for the assembly, genetic engineering and rescue of large DNA viruses such as cytomegalovirus and herpes simplex virus 1^13,14^. While large DNA viruses such as herpesviruses can be rescued by transfecting the entire viral DNA genome into eukaryotic cells, the rescue of RNA viruses requires the transcription of a viral genomic RNA from the cloned DNA template. For coronaviruses that belong to a family of positive-stranded RNA viruses termed *Coronaviridae*, the generation of full-length molecular clones has long been hampered by the extraordinary genome size (27-31 kb) and occasional instability of cloned DNA in *E. coli*. It took until 2000/2001 before unconventional approaches, such as cloning in low copy bacterial artificial chromosomes (BACs) or vaccinia virus, or cloning of subgenomic DNA fragments followed by *in-vitro* ligation, were successful^1-3^. However, each system has caveats leaving the generation of recombinant coronaviruses cumbersome. We therefore aimed to assess the use of the yeast *S. cerevisiae* to assemble and maintain genomes of diverse RNA viruses in order to establish a rapid, stable, and universal reverse genetics pipeline for RNA viruses.

In an effort to adopt a yeast-based reverse genetics platform to RNA viruses we first tested the Murine hepatitis virus strain A59 containing the gene for the green fluorescent protein (MHV-GFP), which is routinely used in our laboratory and has an established vaccinia virus-based reverse genetics platform^15,16^. The overall strategy is shown in Fig. 1a. Viral RNA was prepared from MHV-GFP-infected murine 17Cl-1 cells and used to amplify 7 overlapping DNA fragments by RT-PCR together spanning the MHV-GFP genome from nt 2024 to nt 29672. Fragments containing the 5’- and 3’-termini were PCR-amplified from the vaccinia virus-cloned genome to include a T7-RNA polymerase promoter and a cleavage site (*Pac*I) following the polyA sequence. Overlaps to the TAR vector pVC604 were included in the primers used to amplify the 5’- and 3’-terminal fragments and the pVC604 backbone that was also generated by PCR (Fig. 1b, Table 1, Extended Data Table 1, Extended Data Figure 1a). All DNA fragments were used to transform *Saccharomyces cerevisiae* (strain VL6-48N) and resulting colonies were screened for correct assembly of the YAC containing the MHV genome by multiplex PCRs that covered the junctions of the different fragments. This screen revealed that > 90 % of the clones tested were positive indicating a highly efficient assembly in yeast (Extended Data Figure 1a). To rescue MHV-GFP we randomly chose two clones, purified the YAC, linearized the plasmid using the restriction endonuclease *Pac*I (Extended Data Table 2) and subjected it to T7 RNA polymerase-based *in vitro* transcription to generate capped viral genomic RNA. This RNA was transfected together with an *in vitro* transcribed mRNA encoding the MHV nucleocapsid (N) protein into BHK-MHV-N cells as previously described^15^. Transfected BHK-MHV-N cells were mixed with MHV-susceptible 17Cl-1 cells and cytopathic effect (CPE), virus-induced syncytia, and GFP-expressing cells were readily detectable for both clones within 48 hours, indicating successful recovery of infectious virus (Fig. 1c). Finally, we assessed the replication kinetics of the recovered viruses, which were indistinguishable compared to the parental MHV-GFP (Fig. 1d).

**Table 1:** RNA virus genomes cloned using the synthetic genomics platform

| **Virus** | **Family** | **RNA type** | **Size (kbp)** | **Template** | **Fragment generation** | **Number of assembled fragments\*** | **Virus rescue** |
| --- | --- | --- | --- | --- | --- | --- | --- |
| MHV-GFP | Coronaviridae | (+)RNA | 31.9 | Viral RNA (laboratory virus); DNA clone (vaccinia virus) | RT-PCR/PCR | 9 | yes |
| MERS-CoV | Coronaviridae | (+)RNA | 30.1 | DNA clone (BAC) | PCR | 8 | yes |
| MERS-CoV-GFP | Coronaviridae | (+)RNA | 30.7 | DNA clone (BAC)<br>GFP plasmid DNA | PCR | 10 | yes |
| HCoV-229E | Coronaviridae | (+)RNA | 27.3 | Viral RNA (laboratory virus); DNA clone (vaccinia virus) | RT-PCR/PCR | 13 | in progress |
| HCoV-HKU1 | Coronaviridae | (+)RNA | 29.9 | Synthetic DNA<br>viral RNA (virus isolate) | PCR/RT-PCR | 11 | in progress |
| MERS-CoV-Riyadh-1734-2015 | Coronaviridae | (+)RNA | (30) | Viral RNA (virus isolate) | RT-PCR | 8 | in progress |
| ZIKA virus | Flaviviridae | (+)RNA | 10.8 | Viral RNA (isolate) | RT-PCR | 6 | in progress |
| Human RSV-B | Paramyxoviridae | (-)RNA | 15 | Clinical sample | RT-PCR | 4 | in progress |
| SARS-CoV-2 | Coronaviridae | (+)RNA | 30 | Synthetic DNA/viral RNA | plasmid DNA restriction/ RT-PCR | 12 | yes |
| SARS-CoV-2-GFP | Coronaviridae | (+)RNA | 30.5 | Synthetic DNA/viral RNA | plasmid DNA restriction/RT-PCR/PCR | 14 | yes |

\*excluding the TAR vector fragment

**Figure 1.**
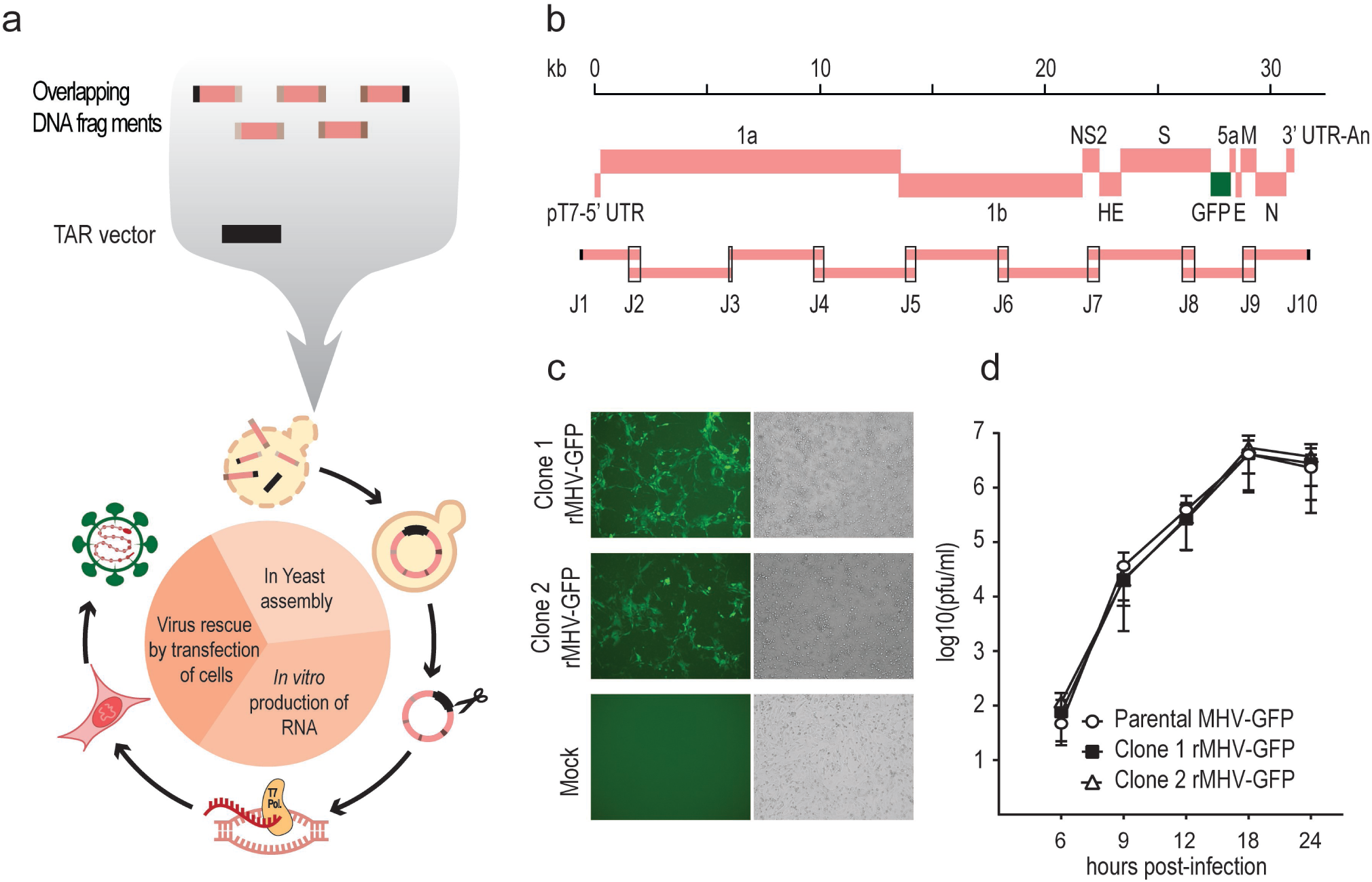
Application of yeast-based TAR cloning to generate viral cDNA clones and the recovery of recombinant MHV-GFP. **a**, Schematic representation of the general workflow of TAR cloning and virus rescue. In-yeast assembly require the delivery of overlapping DNA fragments covering the viral genome and a TAR vector into yeast in one transformation event. Transformed DNA fragments are joined by homologous recombination in yeast generating a YAC-cloned full-length viral genomic cDNA. The *in vitro* production of infectious capped viral RNA starts with the isolation of the YAC, followed by plasmid linearization to provide DNA template for run-off T7 polymerase-based transcription. Virus rescue starts with electroporation of BHK-21 and/or BHK-21 cells expressing the corresponding coronavirus N protein, followed by co-cultivation of electroporated cells with susceptible cell line for virus production and amplification. **b**, Schematic representation of MHV-GFP genome organisation with 9 viral subgenomic overlapping cDNA fragments used for TAR cloning. pT7, T7 RNA polymerase promoter; UTR, untranslated region; An, poly (A) tail; J represents assembly junction between two overlapping DNA fragments. **c**, Recovery of infectious recombinant MHV-GFP. Cell culture supernatants containing viruses produced after virus rescue from two MHV-GFP YAC clones (Clone1, Clone 2) were used to infect 17Cl-1 cells. At 48 hours post-infection, infected cells were visualised for GFP expression (left panels) and by bright field microscopy (right panels). Mock represents 17Cl-1 cells inoculated with supernatant from BHK-MHV-N cells electroporated without viral RNAs. **d**, Replication kinetics of parental MHV-GFP and recombinant MHV-GFP Clone 1 and Clone 2. L929 cells were infected at an MOI of 0.1. Cell culture supernatants were harvested at indicated time points post-infection and titrated by plaque assay. Data are representative of three independent experiments with two replicates per virus in each experiment. Error bars show standard deviation (SD) from the mean. pfu/ml, plaque forming units per millilitre.

To address if the synthetic genomics platform can be applied to other coronaviruses, and if it can be used for rapid mutagenesis, we used a molecular BAC clone of MERS-CoV^17^. We PCR-amplified eight subgenomic overlapping DNA fragments covering the entire MERS-CoV genome (Figure 2a, Extended Data Figure 1b, Extended Data Table 1). The 5’- and 3’- terminal DNA fragments contained the T7-RNA polymerase promoter upstream the MERS-CoV 5’-end and a restriction endonuclease cleavage site *MluI* downstream the polyA sequence, and overlapping sequences with the TAR plasmid pVC604. To mutagenize the MERS-CoV clone and introduce the GFP gene, fragment 7 was modified to overlap with the PCR-amplified GFP gene. Following transformation almost all YAC clones were positive for the junctions indicating a successful assembly (Figure 2a, Extended Data Figure 1b). Virus rescue from cloned DNA was performed as described previously^17^ and virus-induced CPE became apparent in VeroB4 cells inoculated with recovered recombinant MERS-CoV (Fig. 2b). Importantly, we could readily detect green fluorescent VeroB4 cells that had been infected with recovered MERS-CoV-GFP demonstrating that the synthetic genomics platform is suitable to genetically modify coronavirus genomes (Fig. 2b). As expected replication kinetics of recombinant MERS-CoV and MERS-CoV-GFP were slightly reduced compared to cell-culture-adapted MERS-CoV-EMC (Fig. 2c).

**Figure 2.**
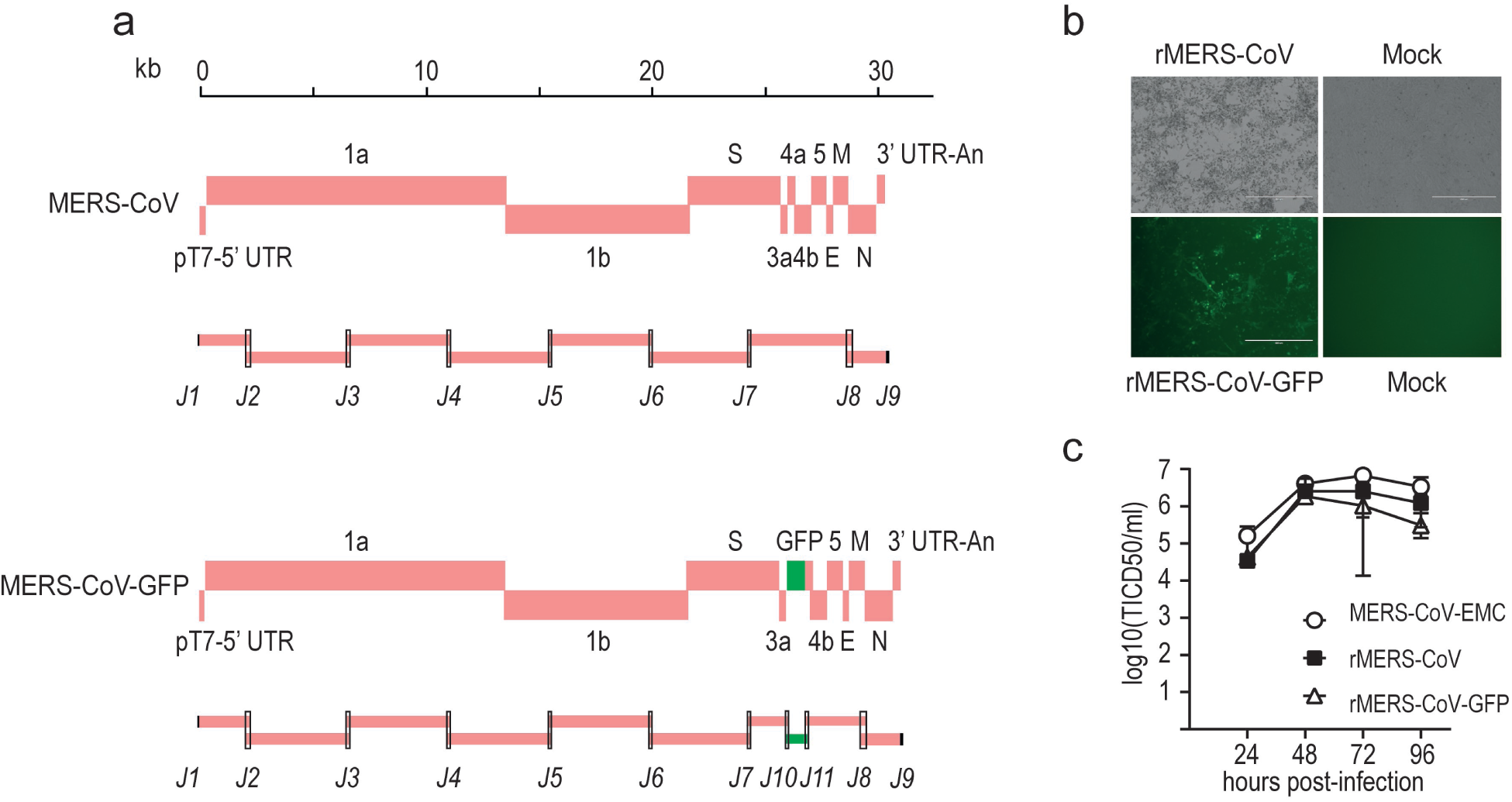
Generation of viral cDNA clones and recovery of recombinant MERS-CoV and MERS-CoV-GFP. **a**, Schematic representation of MERS-CoV (upper panel) and MERS-CoV-GFP (lower panel) genome organization with 8 and 10 viral subgenomic overlapping cDNA fragments, respectively, used for TAR cloning. pT7, T7 RNA polymerase promoter; UTR, untranslated region; An, poly (A) tail; J represents assembly junction between two overlapping DNA fragments. **b**, Rescue of recombinant MERS-CoV and MERS-CoV-GFP. Following delivery of viral RNAs into BHK-21 cells via electroporation the cells were co-cultivated with VeroB4 cells and supernatants containing recombinant viruses produced were used to infect new VeroB4 cells. Infected cells were visualised by bright field microscopy for recombinant MERS-CoV (upper panel; 5 days post infection), and for GFP expression for recombinant MERS-CoV-GFP (lower panel, 3 days post infection). Mock represents VeroB4 cells inoculated with supernatant from BHK-21 cells electroporated without viral RNAs. **c**, Replication kinetics of the MERS-CoV-EMC (laboratory adapted isolate), rMERS-CoV and rMERS-CoV-GFP. VeroB4 cells were infected at an MOI of 0.01. Cell culture supernatants were harvested at indicated time points post infection and titrated by TCID50 assay. Data are representative of three independent experiments with two replicates per virus in each experiment. Error bars show standard deviation (SD) from the mean. TCID50/ml, 50% tissue culture infectious dose per millilitre.

Having shown that the TAR cloning approach is robust and reproducible, we aimed to thoroughly evaluate the system concerning stability of cloned sequences, the range of applicability to other virus genomes, and if molecular clones can even be generated from clinical samples. The yeast clones containing YACs encoding MHV-GFP and MERS-CoV were passaged 15-17 times followed by re-sequencing. Results of this experiment were very supportive of the yeast-based platform (Extended Data Table 3), since the genomes could be stably maintained and no mutations arose. We then embarked on cloning a number of other coronaviruses (HCoV-229E^2^, HCoV-HKU1; GenBank: NC_006577, MERS-CoV-Riyadh-1734-2015; GenBank: MN481979) and viruses of other families, such as ZIKV (family *Flaviviridae*, GenBank: KX377337) and human respiratory-syncytial-virus (hRSV; family *Paramyxoviridae*) (Table 1) that are known to be difficult to clone and stably maintain in *E. coli*. As shown in Extended data Figure 1d-h, we followed the same strategy to prepare the genome termini in cloned plasmids to include genetic elements that are later needed for virus rescue (e.g. T7-RNA polymerase promoter, cleavages sites or ribozyme elements) and produced a set of overlapping DNA fragments covering the rest of the genome. This strategy proved to be successful irrespectively of the source of virus or nucleic acid template or the number of DNA fragments that were used for the one-step assembly in yeast. Of note, we were also able to clone hRSV-B, without any prior information on the virus genotype, directly from a clinical sample (nasopharyngeal aspirate) by designing RSV consensus primers to amplify 4 overlapping DNA fragments (sequence submitted to GenBank). Collectively, these results demonstrate the synthetic genomics platform provides the technical advance to rapidly generate molecular clones of diverse RNA viruses by using viral isolates, cloned DNA, synthetic DNA, and even clinical samples as starting material.

The appearance of a novel CoV in China at the end of 2019 prompted us to go a step further and to test the applicability of our synthetic genomics platform to reconstruct the virus based on its released genome sequence and chemically synthesized DNA fragments. Notably, at the time when the first genome sequences of SARS-CoV-2 were released (Jan 10/11^th^, 2020) no virus isolate has been made available and it took until the end of January 2020 when successful isolation of SARS-CoV-2 was reported from patients in Australia (https://www.the-scientist.com/news-opinion/australian-lab-cultures-new-coronavirus-as-infections-climb-67031). We fragmented the genome into 12 subgenomic DNA fragments ranging in size between 0.5-3.4 kbp (Fig. 3a, Extended Data Figure 1i, Extended Data Tables 1, 4). In parallel, we also aimed to generate a SARS-CoV-2 expressing GFP that could be valuable to facilitate the establishment of serological diagnostics (e.g. virus neutralization assay) and detection in cell culture. To do this we divided fragment 11 into 3 sub-fragments (Fig. 3a, Extended Data Figure 1i, Extended Data Tables 1, 4) to include the GFP sequence, which was inserted in frame within the ORF7a. We noticed at the 5’ end of the reported SARS-CoV-2 sequence that nucleotides 3-5 (5’-AU<u>UAA</u>AGG; Genbank MN996528.1) were different compared to the closely related SARS-CoV (5’-AU<u>AUU</u>AGG; GenBank AY291315) and to the even more closely related bat SARS-related CoVs ZXC21 and ZC45 (5’-AU<u>AUU</u>AGG)^18^, and that for all three the 5’-end has been experimentally determined by 5’-RACE^4,18,19^. Therefore, we designed three 5’-end versions of fragment 1 containing the reported SARS-CoV-2 sequence (fragment 1.3; 5’-AU<u>UAA</u>AGG), a version modified by 3 nucleotides (fragment 1.1; 5’-AU<u>AUU</u>AGG), and a version containing the 124 5’-terminal nucleotides spanning the first four 5’-terminal stem-loop structures of SARS-CoV (fragment 1.2; Fig. 3b). Notably, nucleotide differences between SARS-CoV-2 and SARS-CoV within this region are in agreement with the predicted RNA secondary structures (Fig. 3b).

**Figure 3.**
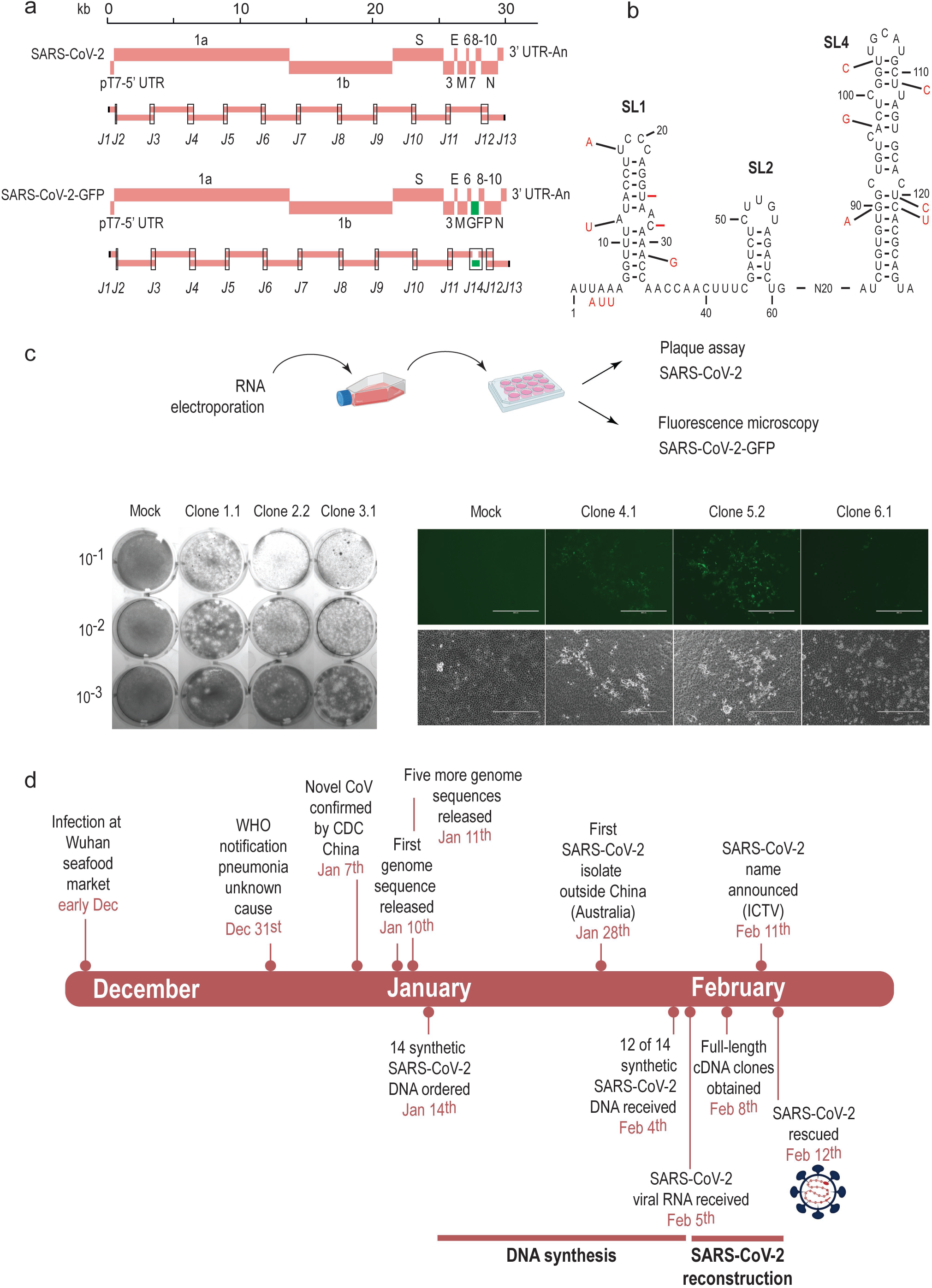
Generation of viral cDNA clones and recovery of recombinant SARS-CoV-2 and SARS-CoV-2-GFP. **a**, Schematic representation of SARS-CoV-2 (upper panel) and SARS-CoV-2-GFP (lower panel) genome organisation with 12 and 14 viral subgenomic overlapping cDNA fragments, respectively, used for TAR cloning. pT7, T7 RNA polymerase promoter; UTR, untranslated region; An, poly (A) tail; J represents assembly junction between two overlapping DNA fragments. **b**, Representation of predicted RNA stem-loop (SL) secondary structures formed in the 5’-UTR of SARS-CoV-2. The SARS-CoV-2 RNA secondary structures were manually adjusted based on RNA structure predictions by Chan et al.^22^. Black letters and numbering represents the SARS-CoV-2 5’-terminal sequence. Red letters depict nucleotides that are different within the SARS-CoV 5’-terminal sequence (“-” indicates nucleotide deletion in SARS-CoV versus SARS-CoV-2). N20 indicates 20 nucleotides. SL structures 1, 2, and 4 are shown according to Chan *et al.*^22^. Note that recombinant virus construct 1 (rSARS-CoV-2 clone 1) and construct 4 (rSARS-CoV-GFP clone 4) contain the SARS-CoV-2 5’-terminus with nucleotides 3-5 (UAA) exchanged by nucleotides 3-5 (AUU) of SARS-CoV; recombinant virus construct 2 (rSARS-CoV-2 clone 2) and construct 5 (rSARS-CoV-2-GFP clone 5) contain a 5’-terminus with the first 124 nucleotides from SARS-CoV; and recombinant virus construct 3 (rSARS-CoV-2 clone 3) and construct 6 (rSARS-CoV-GFP clone 6) contain the authentic SARS-CoV-2 5’-terminus. **c**, Rescue of recombinant rSARS-CoV-2 and rSARS-CoV-2-GFP. The experimental setting is illustrated in the upper panel. Two clones of each construct 1-6 were used to prepare YAC DNAs and *in-vitro* transcribed viral genome RNAs were electroporated into BHK-21 cells and/or BHK-SARS-N together with an mRNA encoding the SARS-CoV-2 N protein. Electroporated cells were co-cultivated with VeroE6 cells in a T25 flask and cell culture supernatants were transferred to 12-well plates to perform plaque assay for rSARS-CoV-2 constructs 1-3 and fluorescence microscopy for rSARS-CoV-2-GFP constructs 4-6. Plaque assays are shown in the lower panel for SARS-CoV-2 clones 1.1, 2.2, and 3.1. VeroE6 cells were infected with 10^−1^, 10^−2^, and 10^−3^ dilutions of 1 ml supernatant from the individual rescue experiments and are compared to non-infected cells (Mock). The expression of GFP in SARS-CoV-2-GFP infected VeroE6 cells is shown for clones 4.1, 5.2, and 6.1. Mock represents non-infected VeroE6 cells. **d**, Summary of the timeline of events of the SARS-CoV-2 outbreak and the work leading to the reconstruction and recovery of rSARS-CoV-2.

We ordered the synthetic DNA constructs on January 14^th^ (Fig. 3d, Extended data Table 4, Extended Data Document 1), and until February 4^th^ 12 of 14 constructs were delivered as sequence-confirmed plasmids. However, fragments 5 and 7 turned out to be problematic to clone in *E. coli* and could not be delivered. Since we received at the same time SARS-CoV-2 viral RNA obtained from an isolate of a Munich patient (SARS-CoV-2/München-1.1/2020/929), we decided to amplify the regions of fragments 5 and 7 by RT-PCR. TAR cloning was immediately initiated, and for all 6 SARS-CoV-2 constructs we obtained correctly assembled molecular clones (3 versions of the 5’-end and each version with and without GFP; Extended Data Fig. 2). Since sequence verification was not possible within this short time frame, we randomly selected two clones for each construct for rescuing recombinant rSARS-CoV-2 and rSARS-CoV-2-GFP. YAC DNA were isolated, *in vitro* transcribed and the resulting RNAs electroporated together with an mRNA encoding the SARS-CoV-2 N protein into BHK-21 and, in parallel, into BHK-SARS-N cells expressing the SARS-CoV N protein^20^. The BHK-21 and BHK-SARS-N cells were seeded over VeroE6 cells and at two days post electroporation we observed green fluorescent signals in cells that received the GFP-encoding SARS-CoV-2 RNAs. Indeed supernatants transferred to fresh VeroE6 cells contained infectious recombinant viruses for almost all recombinant rSARS-CoV-2 and rSARS-CoV-GFP constructs (Extended Data Fig. 2). As shown in Figure 3c for SARS-CoV-2 clones 1.1, 2.2, and 3.1, plaques were detectable on VeroE6 cells that were inoculated with supernatants containing the RNA-electroporated cells, demonstrating the infectious virus has been recovered for recombinant rSARS-CoV-2 and for the variants containing modified 5’-termini. Similarly, we could detect green fluorescent cells on VeroE6 monolayers that were inoculated with supernatants from RNA-electroporated cells of rSARS-CoV-2-GFP clones 4.1, 5.2 and 6.1, demonstrating that recombinant GFP-expressing viruses have been recoverd for all 5’-end variants. This result demonstrates the full functionality of the SARS-CoV-2 reverse genetics system and also shows that the 5’-ends of SARS-CoV and bat SARS-related CoVs ZXC21 and ZC45 are compatible with the replication machinery of SARS-CoV-2.

We expect that the fast, robust and versatile synthetic genomics platform we describe here will provide new insights into the molecular biology and pathogenesis of a number of emerging RNA viruses and will specifically facilitate the molecular characterization of the novel coronavirus SARS-CoV-2. Although homologous recombination in yeast has already been used for the generation of a number of molecular virus clones in the past^13,14,21^, we present here a thorough evaluation of the feasibility of this approach to rapidly generate full-length cDNAs for large RNA viruses that have a known history of instability in *E. coli*. We show that one main advantage of the TAR cloning system is that the viral genomes can be fragmented to at least 14 overlapping fragments and re-assembled with remarkable efficacy (usually >90% of the clones are correct). This allowed us to complete the cloning and rescue of rSARS-CoV-2 and rSARS-CoV-2-GFP within one week. It should also be noted that we see considerable potential to reduce the time of DNA synthesis. Currently, synthetic DNA fragments are still routinely cloned in *E. coli*, which turned out to be problematic for SARS-CoV-2 fragments 5 and 7. We could however use 4 and 3 shorter dsDNA parts of fragments 5 and 7, respectively, that we received, to assemble the full-length fragments 5 and 7 by TAR cloning which is an additional proof of the superior cloning efficiency of our yeast system versus *E. coli*-based systems. This allowed us to generate a molecular clone of SARS-CoV-2 by using exclusively chemically synthesized DNA, and the rescue of this version is currently in progress.

The appearance of SARS-CoV-2 emphasizes the need for preparedness to rapidly respond to emerging virus threats. While the availability of SARS-CoV-2 isolates is still limited, full virus genome sequences were disclosed very early during the outbreak. The rapidity of our synthetic genomics approach to generate SARS-CoV-2 and the applicability to other emerging RNA viruses make this system an attractive alternative to provide infectious virus to health authorities and diagnostic laboratories without the need to have access to clinical samples. Finally, as the SARS-CoV-2 outbreak is ongoing, we expect to see sequence variations and possibly phenotypic changes of evolving SARS-CoV-2 in the human host. With the synthetic genomics platform it is now possible to introduce such sequence variations into the infectious clone and to functionally characterize SARS-CoV-2 evolution in real-time.

## Acknowledgements

This work was supported by the European Commission (Marie Sklodowska-Curie Innovative Training Network “HONOURS”; grant agreement No 721367), the Swiss National Science Foundation (SNF; grants CRSII3_160780, and 310030_173085), the German Research Council (DFG; grants SFB-TR84 (TRR 84/3, A07), and DR 772/7-2), the Federal Ministry of Education and Research (BMBF; grant RAPID, #01KI1723A) and by core funds of the University of Bern. We thank the J. Craig Venter Institute (JCVI) for provision of the TAR vectors. We thank specifically Eric Sun and Tibbers Zhang and are grateful for the timely provision of the synthetic DNA fragments from GenScript. We thank Jeanne Peters Zocher from VETCOM Zürich for her advice in generating the figures and we thank Franziska Suter-Riniker and Pascal Bittel, Institute for Infectious Diseases, University of Bern, for providing clinical samples. We thank Philippe Plattet, Markus Gerber, Martina Friesland, Marco Alves, Nathalie Vielle, Beatrice Zumkehr, Melanie Brügger, Daniel Brechbühl for reagents, technical advise and helpful discussions.

## Authors contributions

VT and JJ conceived the study. TT, NE and FL performed most of the experiments. PV, HS, JP, SSt, MH, AK, MG, LL, LH, MW, SP, DH, VC, SC, SSch, DM, DN, MM, CD, RD, did experimental work and/or provided essential experimental systems and reagents. TT, NE, FL, HS, JK, and RD performed sequencing including computational analyses. VT, JJ, TT, FL, NE wrote the manuscript and made the figures. All authors read and approved the final manuscript.

## Ethical statement

The authors are aware that this work contains aspects of Dual Use Research of Concern (DURC). The benefits were carefully balanced against the risks and the benefits outweigh the risks. Permission to generate and work with recombinant SARS-CoV-2 and SARS-CoV-2-GFP was granted by the Swiss Federal Office of Public Health (#A131191/3) with consultation of the Federal Office for Environment, Federal Food Safety and Veterinary Office, and the Swiss Expert Committee for Biosafety.

## Additional information

Correspondence and requests for material should be addressed to V.T. or J.J.

## Figure Legends

**Extended Data Figure 1**

**a**, Schematic representation of the MHV-GFP genome organisation (upper panel) and nine viral subgenomic cDNA fragments (F1-9) used for TAR cloning. Viral open reading frames (ORFs), the ORF for GFP and sequence elements at the 5’- and 3’-untranslated regions (UTRs) are indicated. Primers used to generate the fragments are listed in Extended Data Table 1. J2-9 represent the junctions, i.e. overlapping regions, between the subgenomic cDNA fragments. J1 and J10 represent junctions with the TAR vector. Gel images show results from two multiplex PCRs that were designed to detect the presence of correctly recombined junctions J1-10, confirming the proper assembly of the YAC containing the viral genome in 12 out of 12 clones. kb, kilobases; pT7, T7 RNA polymerase promoter; An, poly (A) tail.

**b**, Schematic representation of the MERS-CoV genome organisation (upper panel) and eight viral subgenomic cDNA fragments (F1-8) used for TAR cloning. Viral open reading frames (ORFs) and sequence elements at the 5’- and 3’-untranslated regions (UTRs) are indicated. Primers used to generate the fragments are listed in Extended Data Table 1. J2-8 represent the junctions, i.e. overlapping regions, between the subgenomic cDNA fragments. J1 and J9 represent junctions with the TAR vector. Gel image shows results from a multiplex PCR that was designed to detect the presence of correctly recombined junctions J1-9, confirming the proper assembly of the YAC containing the viral genome in 6 out of 8 clones. kb, kilobases; pT7, T7 RNA polymerase promoter; An, poly (A) tail.

**c**, Schematic representation of the MERS-CoV-GFP genome organisation (upper panel) and ten viral subgenomic cDNA fragments (F1-10 and F13) used for TAR cloning. Viral open reading frames (ORFs), ORF for GFP and sequence elements at the 5’- and 3’-untranslated regions (UTRs) are indicated. Primers used to generate the fragments are listed in Extended Data Table 1. J2-8 and J10-11 represent the junctions, i.e. overlapping regions, between the subgenomic cDNA fragments. J1 and J9 represent junctions with the TAR vector. Gel images show results from two multiplex PCRs that were designed to detect the presence of correctly recombined junctions J1-11, confirming the proper assembly of the YAC containing the viral genome in 5 out of 8 clones. kb, kilobases; pT7, T7 RNA polymerase promoter; An, poly (A) tail.

**d**, Schematic representation of the HCoV-229E genome organisation (upper panel) and thirteen viral subgenomic cDNA fragments (F1-13) used for TAR cloning. Viral open reading frames (ORFs) and sequence elements at the 5’- and 3’-untranslated regions (UTRs) are indicated. Primers used to generate the fragments are listed in Extended Data Table 1. J2-13 represent the junctions, i.e. overlapping regions, between the subgenomic cDNA fragments. J1 and J14 represent junctions with the TAR vector. Gel images show results from two multiplex PCRs that were designed to detect the presence of correctly recombined junctions J1-14, confirming the proper assembly of the YAC containing the viral genome in 7 out of 10 clones. kb, kilobases; pT7, T7 RNA polymerase promoter; An, poly (A) tail.

**d**, Schematic representation of the HCoV-HKU1 genome organisation (upper panel) and eleven viral subgenomic cDNA fragments (F1-11) used for TAR cloning. Viral open reading frames (ORFs) and sequence elements at the 5’- and 3’-untranslated regions (UTRs) are indicated. Primers used to generate the fragments are listed in Extended Data Table 1. J2-11 represent the junctions, i.e. overlapping regions, between the subgenomic cDNA fragments. J1 and J12 represent junctions with the TAR vector. Gel images show results from two multiplex PCRs that were designed to detect the presence of correctly recombined junctions J1-12, confirming the proper assembly of the YAC containing the viral genome in 14 out of 14 clones. kb, kilobases; pT7, T7 RNA polymerase promoter; An, poly (A) tail.

**e**, Schematic representation of the MERS-CoV-Riadh-1734-2015 genome organisation (upper panel) and eight viral subgenomic cDNA fragments (F1-8) used for TAR cloning. Viral open reading frames (ORFs) and sequence elements at the 5’- and 3’-untranslated regions (UTRs) are indicated. Primers used to generate the fragments are listed in Extended Data Table 1. J2-8 represent the junctions, i.e. overlapping regions, between the subgenomic cDNA fragments. J1 and J9 represent junctions with the TAR vector. Gel images show results from a multiplex PCR and a simplex PCR, both of which were designed to detect the presence of correctly recombined junctions J1-9, confirming the proper assembly of the YAC containing the viral genome in 5 out of 8 clones. kb, kilobases; pT7, T7 RNA polymerase promoter; An, poly (A) tail.

**f**, Schematic representation of the ZIKA virus genome organisation (upper panel) and six viral subgenomic cDNA fragments (F1-6) used for TAR cloning. Viral open reading frames (ORFs) and sequence elements at the 5’- and 3’-untranslated regions (UTRs) are indicated. Primers used to generate the fragments are listed in Extended Data Table 1. J2-5 represent the junctions, i.e. overlapping regions, between the subgenomic cDNA fragments. J1 and J6 represent junctions with the TAR vector. Gel image shows results from a multiplex PCR that was designed to detect the presence of correctly recombined junctions J1-6, confirming the proper assembly of the YAC containing the viral genome in 3 out of 15 clones. kb, kilobases; pT7, T7 RNA polymerase promoter; HDR, hepatitis delta virus ribozyme.

**g**, Schematic representation of the human RSV-B virus genome organisation (upper panel) and six viral subgenomic cDNA fragments (F1-6) used for TAR cloning. Viral open reading frames (ORFs) and sequence elements at the 5’- and 3’-untranslated regions (UTRs) are indicated. Primers used to generate the fragments are listed in Extended Data Table 1. J1-5 represent the junctions, i.e. overlapping regions, between the subgenomic cDNA fragments. J1 and J5 contain junctions with the TAR vector. Gel image shows results from a multiplex PCR that was designed to detect the presence of correctly recombined junctions J1-5, confirming the proper assembly of the YAC containing the viral genome in 7 out of 8 clones. kb, kilobases; pT7, T7 RNA polymerase promoter; HHrb, hammerhead ribozyme, Rb-T7ter, Ribozyme and T7 RNA polymerase terminator seqeunce.

**h**, Schematic representation of the SARS-CoV-2 genome organisation (upper panel) and twelve viral subgenomic cDNA fragments (F1-12) used for TAR cloning. Viral open reading frames (ORFs) and sequence elements at the 5’- and 3’-untranslated regions (UTRs) are indicated. Primers used to generate the fragments are listed in Extended Data Table 1. J2-12 represent the junctions, i.e. overlapping regions, between the subgenomic cDNA fragments. J1 and J13 represent junctions with the TAR vector. Gel images show results from two multiplex PCRs that were designed to detect the presence of correctly recombined junctions J1-13, confirming the proper assembly of the YAC containing the viral genome in 6 out of 6 clones. kb, kilobases; pT7, T7 RNA polymerase promoter; An, poly (A) tail.

**i**, Schematic representation of the SARS-CoV-2 genome organisation (upper panel) and fourteen viral subgenomic cDNA fragments (F1-10 and F12-15) used for TAR cloning. Viral open reading frames (ORFs), ORF for GFP and sequence elements at the 5’- and 3’-untranslated regions (UTRs) are indicated. Primers used to generate the fragments are listed in Extended Data Table 1. J2-12 and J14 represent the junctions, i.e. overlapping regions, between the subgenomic cDNA fragments. J1 and J13 represent junctions with the TAR vector. Gel images show results from two multiplex PCRs and a simplex PCR, all of which were designed to detect the presence of correctly recombined junctions J1-13 and J14 respectively, confirming the proper assembly of the YAC containing the viral genome in 6 out of 6 clones. kb, kilobases; pT7, T7 RNA polymerase promoter; An, poly (A) tail.

**Extended Data Figure 2**

**a**, Six constructs were initially designed based on 3 different 5’-UTR regions. These regions contain i) a modified sequence of the 5’-UTR region of the SARS-CoV-2 (5’-AT<u>AUU</u>AGG; 5’-terminal 8 nucleotides are shown, modified nucleotides underlined) were nucleotides 3-5 (UAA) of SARS-CoV-2 were changed to AUU to match the SARS-CoV nucleotides 3-5 (synthetic fragment 1.1 for constructs 1 and 4), ii) a SARS-CoV-2 5’-terminus with the first 124 nucleotides exchanged by the corresponding 5’-terminal sequence of SARS-CoV (synthetic fragment 1.2; construct 2 and 5) and iii) the reported sequence of the SARS-CoV-2 (GISAID: EPI_ISL_402119) (synthetic fragment 1.3; constructs 3 and 6). After yeast transformation, 10 colonies were randomly picked for each of the six constructs and all the junctions bridging the overlapping fragments were verified by multiplex PCR. For each construct, two clones (x.1 and x.2) were randomly selected and YAC DNAs were isolated (12 clones in total). RNAs were generated by *in vitro* transcription and electroporated together with an mRNA encoding the SARS-CoV-2 N protein either into BHK-21 cells (12 clones) or BHK-SARS-N cells (cells expressing the SARS-CoV N protein) (6 clones). Electroporated cells were then co-cultivated with susceptible VeroE6 cells to rescue the recombinant viruses. P.0 supernatants were collected at different time points after electroporation (from 2 dpe to 5 dpe) and transferred to VeroE6 cells to generate P.1 virus stocks, and in parallel to demonstrate presence of infectious virus by plaque assay (virus clones that do not encode GFP) or fluorescence microscopy (GFP-encoding virus clones). YAC, yeast artificial chromosome; h.p.e, hours post-electroporation; d.p.e, days post-electroporation; CPE, cytopathic effects; GFP, green fluorescent protein; pfu/ml, plaque forming units per millilitre.

