## Extended Data Methods for "Rapid reconstruction of SARS-CoV-2 using a synthetic genomics platform"

### Cells and general culture conditions

Vero, VeroB4 and VeroB6 cells were cultured in Dulbecco’s modified Eagle’s medium (DMEM). BHK-21, BHK-SARS-N (BHK-21 cells expressing SARS N protein)^1^, L929 and murine 17Cl-1 cells were grown in minimal essential medium (MEM). Both types of media were supplemented with 10% fetal bovine serum, 1% non-essential amino acids and 1% penicillin/streptomycin. All cells were maintained at 37^o^C and in 5% CO_2_ atmosphere.

### Cultured viruses

In our lab, MHV-GFP^2,3^ and HCoV-229E^4^ were cultured in murine 17Cl-1 and Huh-7 cells, respectively. MERS-CoV-EMC^5^ was cultured in VeroB4 cells. HCoV-HKU1 strain Caen-1 (GenBank: NC_006577) was an isolate and cultured on human airway epithelia cultures^6^. ZIKV strain PRVABC-59 (GenBank: KX377337) was provided by M. Alves (Institute of Virology and Immunology, Bern, Switzerland) and cultured on Vero cells.

### Bacterial and yeast strains

*Escherichia coli* DH5α (Thermo Scientific™) and TransforMax™ Epi300™ electrocompetent cells (Epicentre) strains were used to propagate the pVC604 and pCC1BAC-His3 TAR vectors, respectively. The bacteria were grown in lysogeny media (LB) supplemented with appropriate antibiotics at 37°C overnight.

*E. coli* Epi300™ cells transformed with the different SARS-CoV-2 synthetic fragments cloned in pUC57/pUC57mini were grown at 30°C to lower instability/toxicity risks.

*Saccharomyces cerevisiae* VL6-48N (MATα trp1-Δ1 ura3-Δ1 ade2-101 his3-Δ200 lys2 met14 cir°) was used for all yeast transformation experiments^7^. Yeast cells were first grown in YPDA broth (Takara Bio), and transformed cells were plated on minimal synthetic defined (SD) agar media without histidine (SD-His) (Takara Bio).

**Viral RNA, infectious cDNA clones and synthetic DNA fragments (Table 1)**

In general, viral cDNA fragments were obtained via RT-PCR of viral RNA extracted from cultured viruses or isolates and from clinical sample, using SuperScript™ IV One-Step RT-PCR System following manufacturer’s instructions. Additionally, some fragments were amplified by PCR from vaccinia virus-cloned cDNA, BAC-cloned cDNA and plasmid-cloned synthetic DNA, using CloneAmp™ HiFi PCR Premix according to manufacturer’s instructions. Accessory sequences, i.e. enhanced GFP and porcine teschovirus-1 2A (P2A) for MERS-CoV-GFP construct, TurboGFP for SARS-CoV-2-GFP, and T7 RNA polymerase promoter-hammerhead ribozyme and ribozyme-T7 terminator for human RSV-B, were amplified from plasmids kindly provided by collaborators.

For all coronaviruses, the first and last fragments for TAR cloning contain T7 RNA polymerase promoter sequence upstream of the viral 5’-UTR, and a poly(A) tail downstream of the 3’-UTR followed by an unique restriction site (Extended Data Table 2), respectively.

Four HCoV-HKU1 synthetic fragments, i.e. fragment 1 to 4, were provided individually in pUC57 by GenScript. For MERS-CoV-Riyadh-1734-2015 (GenBank: MN481979), fragment 1 and 8 were chemically synthesized and cloned in pUC57 by GenScript. Both of these fragments contain homologous regions to TAR vectors pVC604 and pCC1BAC-His3. Similarly, synthetic and pUC57-cloned ZIKA virus fragment 6 contains a hepatitis delta virus ribozyme sequence and pCC1BAC-his3 homology downstream of the viral 3’-UTR.

For SARS-CoV-2, synthetic DNA fragments were delivered cloned in pUC57 or pUC57mini by GenScript (Extended Data Document 1, Extended Data Table 4). Fragment 1.1, 1.2, 1.3 and fragment 12 possess homologous sequences to pCC1BAC-his3. Each fragment was sequence verified using Sanger sequencing after plasmid isolation using Qiagen® Midiprep kit (Qiagen). Fragments were released from the vector using the restriction enzymes provided in Extended Data Table 2. Synthetic fragments were subsequently gel purified using standard methods^8^. DNA concentrations and purities of all fragments to be used for TAR cloning were determined using NanoDrop™ 2000/2000c Spectrophotometer (Thermo Scientific™).

### In-yeast cloning of viral genomes using Transformation-Associated Recombination (TAR) strategy

The vectors pVC604^9^ and pCC1BAC-His3^10^ were used for TAR cloning. These vectors were amplified by PCR using primers containing at least 45-bp overlaps matching the 5’ and 3’-ends of different viral genomes (Extended Data Table 1). Amplification was performed using KOD Hot Start DNA polymerase (Merck Millipore) according to manufacturer’s instruction. Templates used for generating fragments to be assembled in *S. cerevisiae* are displayed in Table 1 and were generated as described above.

Yeast transformation was done with the high efficiency lithium acetate/SS carrier DNA/PEG method described previously ^11^. Briefly, yeast cells were grown in rich YPDA medium (Takara Bio) at 30°C under agitation until an OD_600nm_ of 1.0 was reached. Three ml of yeast culture were used per transformation event. DNA mixtures were prepared beforehand and contained 100-200 fmol of 3’/5’ open ends for all fragments. Transformation mixtures were plated onto SD-His plates (Takara Bio) and incubated at 30°C for 48 hours. Colonies were resuspended in 20 µl of SD-His broth, which was subsequently used to screen for positive clones. DNA extraction was performed following the GC prep method ^12^. Colonies were screened by multiplex PCR using the Qiagen® Multiplex PCR kit (Qiagen) according to the manufacturer’s instruction. One or two multiplex PCRs encompassing different subsets of primer pairs, covering all recombination junctions, were used for the screening process (Extended Data Table 1). Clones tested positive for all assembly junctions were grown in SD-His until late logarithmic phase, and plasmids were extracted from 500 ml culture using the QIAGEN® Maxiprep Kit (QIAGEN, Germany) with modifications. Briefly, 10 ml of Buffer P1 was supplemented with 1 ml of zymolyase solution (10mg/mL Zymolyase 100-T; 50mM Tris-HCl pH7,5; 50% (v/v) glycerol) and 100µl of β-mercapthoethanol. The mixture was incubated for 1 hour at 37°C prior to the addition of buffer P2. The rest of the protocol followed manufacturer’s instructions.

### Stability testing of the YAC containing entire RNA virus genomes in yeast

Stability of the viral genomes maintained as YAC in *S. cerevisiae* was tested for the clones containing MHV-GFP or MERS-CoV over one week. A single colony was grown in SD-His in a 20 ml culture, 1 ml aliquots were removed and expanded in fresh medium every 12 hours. The generation time for each of the clones was estimated to range from 150-160 minutes. After 15-17 passages each YAC clone was isolated and subjected to next generation sequencing (MinION, Oxford Nanopore Technologies).

### Virus rescue

The YAC containing viral cDNA was cleaved at unique restriction site located downstream of the 3’-end poly(A) tail (Extended Data Table 2). Briefly, 1-2 µg of phenol-chloroform extracted and ethanol precipitated DNA resolved in nuclease free water was used for *in vitro* transcription using T7 RiboMAX^TM^ expression large scale RNA production system with m7G(5’)ppp(5’)G cap provided as described previously ^4^. Additionally, a similar protocol was performed on a PCR product of N gene from corresponding coronaviruses, producing a capped mRNA encoding the N protein. 1-10 µg of *in vitro* transcribed viral genome RNA was electroporated together with 2 µg of the N gene transcript into BHK-21 cells and/or BHK-21 cells expressing the corresponding coronavirus N protein. Electroporated cells were co-cultivated with susceptible murine 17Cl-1, VeroB4 and VeroE6 cells to rescue recombinant MHV-GFP, MERS-CoV(-GFP) and SARS-CoV-2(-GFP), respectively. Progeny viruses harvested from immediate post-electroporation supernatant were termed passage 0 viruses, and used to produce stocks for further analysis.

All work involving the rescue and characterisation of recombinant MERS-CoV, SARS-CoV and SARS-CoV-2 was performed in a biosafety level 3 laboratory at the Institute of Virology and Immunology, Mittelhäusern, Switzerland under appropriate safety measures with respect to personal and environmental protection.

### Virus growth kinetics

### For MHV-GFP, twenty-four hours before infection L929 cells were seeded in a 24-well plate at a density of 0.3 x 10^6^ cells/ml. At the time of infection, cells were washed once with PBS and inoculated with viruses. After two hours, virus-containing supernatant was removed, and cells were washed 3 times with PBS and supplied with MEM as described above. Cell culture supernatants were collected at 6, 9, 12, 18 and 24 hours post infection. Similar protocol was applied for MERS-CoV(-GFP) using VeroB4 cells. Cell supernatants were collected at 24, 48, 72 and 96 hours post infection.

### Plaque assay and TCID50.

MHV-GFP plaque forming units (PFU) per millilitre were determined by plaque assay in L929 cells as described previously ^2^. Briefly, twenty-four hours prior to infection L929 cells were seeded in a 24-well plate at a density of 3 x 10^5^ cells/ml. At the time of infection, cells were washed with PBS and inoculated with viruses serially diluted in cell culture medium at 1:10 dilution. Two hours post-inoculation, cells were washed with PBS, and overlaid with 2% methylcellulose mixed at 1:1 with 2X DMEM supplemented with 20% fetal bovine serum and 2% penicillin/streptomycin. After twenty-four hours of incubation, the overlay was removed, and cells were fixed and stained with crystal violet.

The 50% tissue culture infectious dose (TCID50) assay was performed for MERS-CoV(-GFP) in VeroB4 cells. Briefly, VeroB4 cells were seeded 24 hours before infection in 96-wells at a density of 5 x 10^4^ cells/ml. Viruses were serially diluted at 1:10 dilution from 10^-1^ to 10^-7^, one dilution per well and six replicates per virus. After 72 hours of incubation, the media were removed, and cells were fixed and stained with crystal violet. The TCID50 titre was determined using the Spearman-Kaerber method^13^.

For SARS-CoV-2(-GFP), plaque assay determining pfu/ml was performed using VeroE6 cells in a 6-well format. Twenty-four hours before infection, VeroE6 cells were seeded at a density of 3 x 10^5^ cells/well. At the time of infection, cells were washed with PBS and inoculated with viruses serially diluted in cell culture medium at 1:10 dilution. One hour post-inoculation, cells were washed with PBS, and overlaid with 2.4% Avicel mixed at 1:1 with 2X DMEM supplemented with 20% fetal bovine serum and 2% penicillin/streptomycin. After 48 hours of incubation, the overlay was removed, and cells were fixed and stained with crystal violet.

### Sequencing and computational analysis.

For SARS-CoV-2(-GFP), full-length sequences of viral cDNA cloned in pCC1BAC-His3 were confirmed by Sanger sequencing (Microsynth AG, Switzerland). For all other viruses, sequencing of full-length viral genomes cloned in yeast was carried out using the nanopore sequencer MinION from Oxford Nanopore Technologies according to standard protocols. The operating software MinKNOW performed data acquisition and real-time basecalling, generating data as *fast5* and/or *fastq* files. Subsequently, Python command-line *qcat* (Mozilla Public License 2.0. Copyright © 2018 Oxford Nanopore Technologies Ltd. qcat (v1.1.0). Available at: http://www.github.com/nanoporetech/qcat) was run to demultiplex nanopore reads from *fastq* files. Alignment of demultiplexed reads to reference sequences was carried out using *Minimap2* program^14^, producing a *fasta* file. Mutations of consensus sequences were further confirmed by Sanger sequencing. Apart from MinION data handling, other sequence analyses were performed using Geneious Prime ® 2019.2.3. Results from virus growth kinetics were analysed and graphically presented using GraphPad Prism version 8.3.0 for Windows. All figures were created with Adobe Illustrator and Biorender.com.

### Data Availability

The following genome sequences have been submitted to GenBank: rSARS-CoV-2 (#pending), hRSV/B/Bern/2019 (#pending); MERS-CoV-Riyadh-1734-2015 (#MN481979).
