## Extended Data Figure 1 for "Rapid reconstruction of SARS-CoV-2 using a synthetic genomics platform"

a

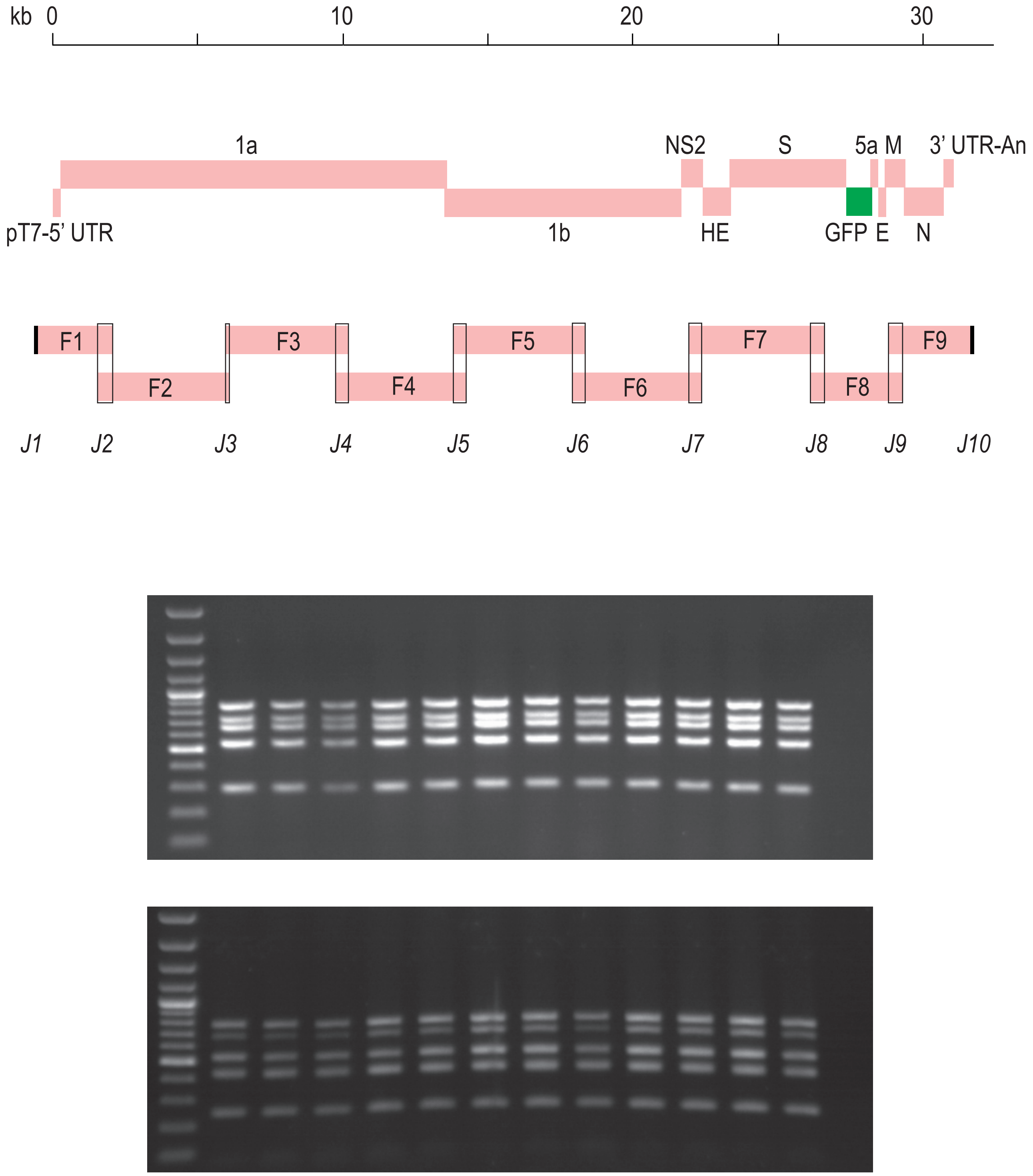


**a,** Schematic representation of the MHV-GFP genome organisation (upper panel) and nine viral subgenomic cDNA fragments (F1-9) used for TAR cloning. Viral open reading frames (ORFs), the ORF for GFP and sequence elements at the 5'- and 3'-untranslated regions (UTRs) are indicated. Primers used to generate the fragments are listed in Extended Data Table 1. J2-9 represent the junctions, i.e. overlapping regions, between the subgenomic cDNA fragments. J1 and J10 represent junctions with the TAR vector. Gel images show results from two multiplex PCRs that were designed to detect the presence of correctly recombined junctions J1-10, confirming the proper assembly of the YAC containing the viral genome in 12 out of 12 clones. kb, kilobases; pT7, T7 RNA polymerase promoter; An, poly (A) tail.

Extended Data Figure 1

b

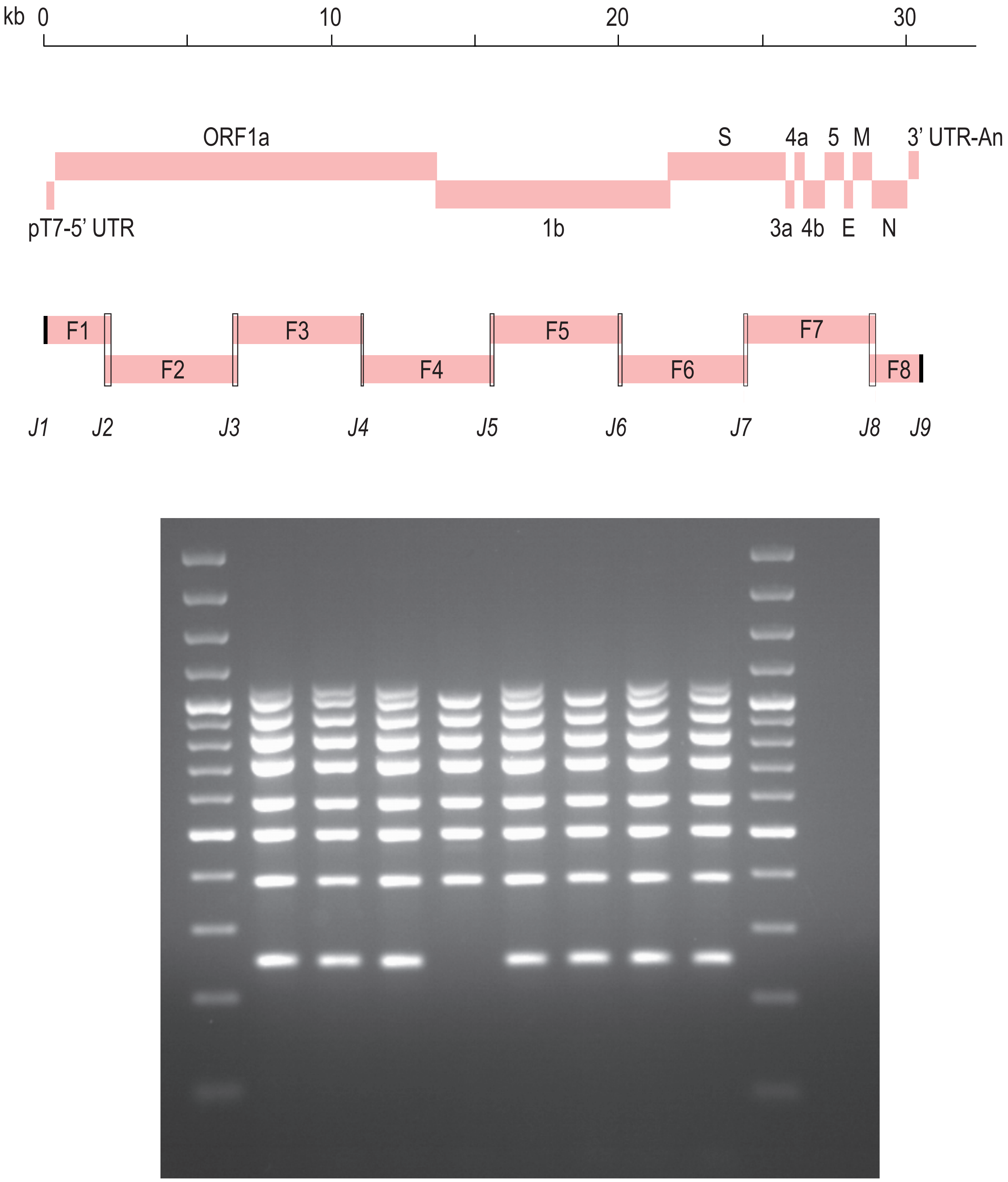


**b,** Schematic representation of the MERS-CoV genome organisation (upper panel) and eight viral subgenomic cDNA fragments (F1-8) used for TAR cloning. Viral open reading frames (ORFs) and sequence elements at the 5'- and 3'-untranslated regions (UTRs) are indicated. Primers used to generate the fragments are listed in Extended Data Table 1. J2-8 represent the junctions, i.e. overlapping regions, between the subgenomic cDNA fragments. J1 and J9 represent junctions with the TAR vector. Gel image shows results from a multiplex PCR that was designed to detect the presence of correctly recombined junctions J1-9, confirming the proper assembly of the YAC containing the viral genome in 6 out of 8 clones. kb, kilobases; pT7, T7 RNA polymerase promoter; An, poly (A) tail.

Extended Data Figure 1

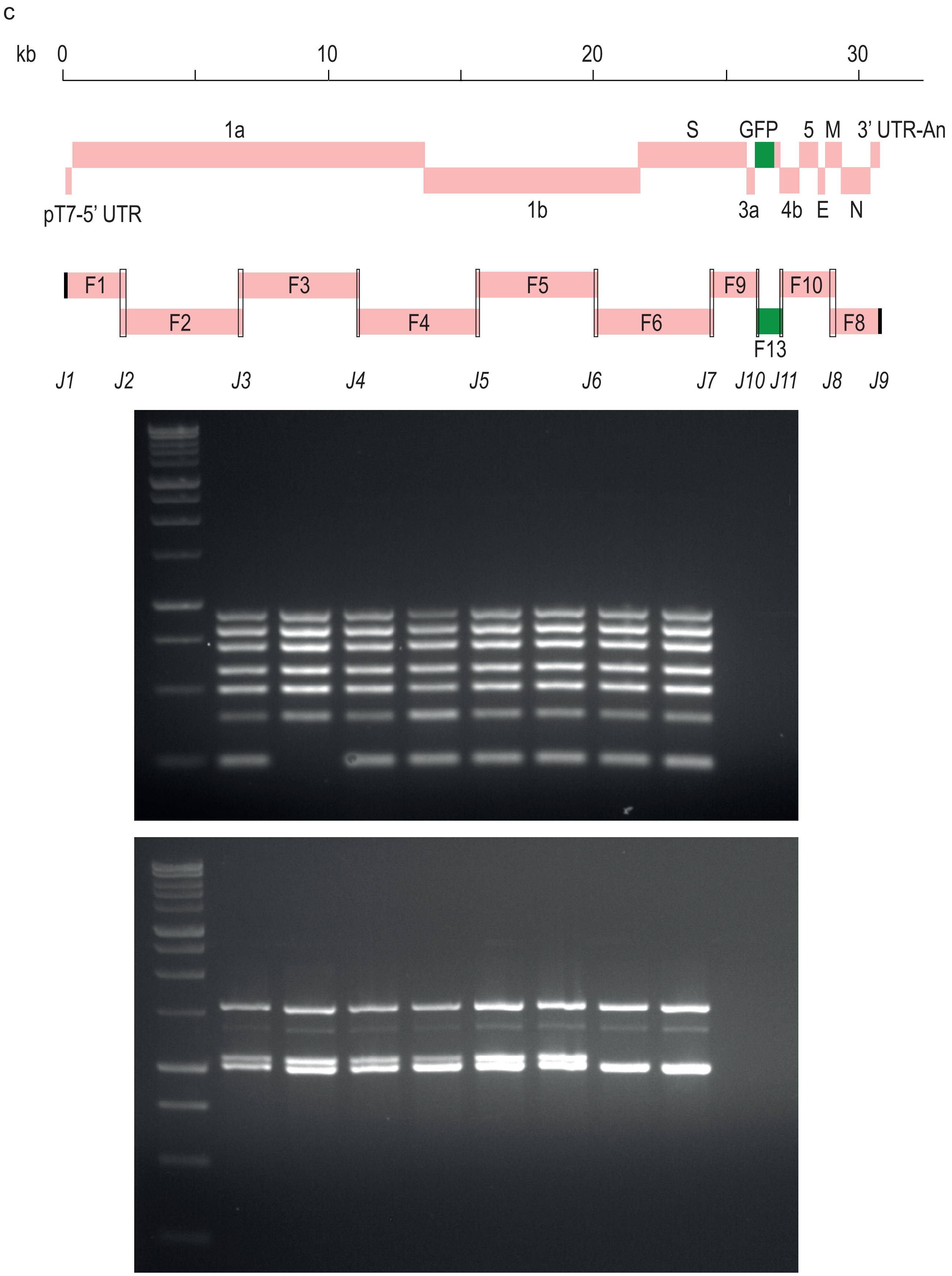


**c,** Schematic representation of the MERS-CoV-GFP genome organisation (upper panel) and ten viral subgenomic cDNA fragments (F1-10 and F13) used for TAR cloning. Viral open reading frames (ORFs), ORF for GFP and sequence elements at the 5'- and 3'-untranslated regions (UTRs) are indicated. Primers used to generate the fragments are listed in Extended Data Table 1. J2-8 and J10-11 represent the junctions, i.e. overlapping regions, between the subgenomic cDNA fragments. J1 and J9 represent junctions with the TAR vector. Gel images show results from two multiplex PCRs that were designed to detect the presence of correctly recombined junctions J1-11, confirming the proper assembly of the YAC containing the viral genome in 5 out of 8 clones. kb, kilobases; pT7, T7 RNA polymerase promoter; An, poly (A) tail.

Extended Data Figure 1

d

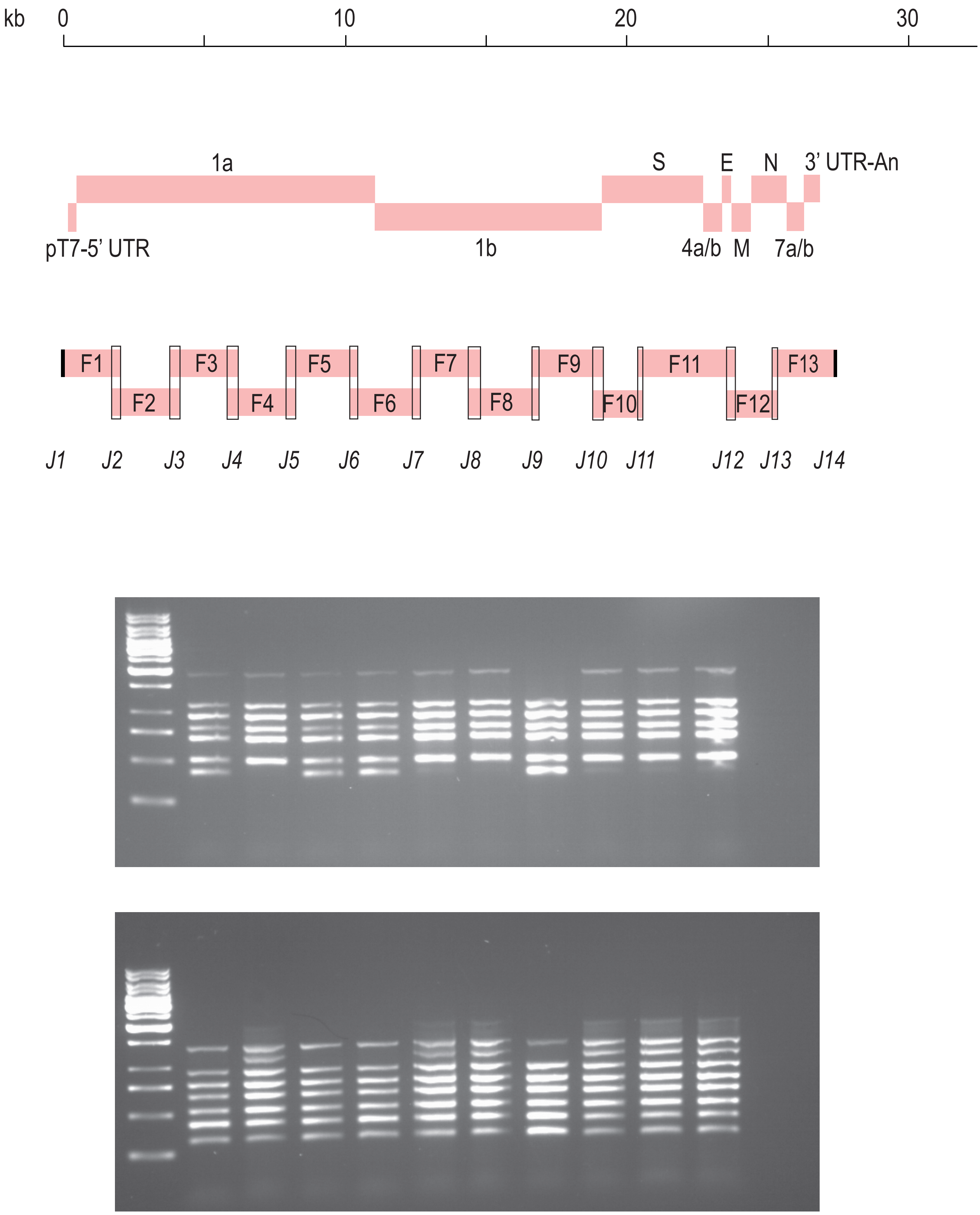


**d,** Schematic representation of the HCoV-229E genome organisation (upper panel) and thirteen viral subgenomic cDNA fragments (F1-13) used for TAR cloning. Viral open reading frames (ORFs) and sequence elements at the 5’- and 3’-untranslated regions (UTRs) are indicated. Primers used to generate the fragments are listed in Extended Data Table 1. J2-13 represent the junctions, i.e. overlapping regions, between the subgenomic cDNA fragments. J1 and J14 represent junctions with the TAR vector. Gel images show results from two multiplex PCRs that were designed to detect the presence of correctly recombined junctions J1-14, confirming the proper assembly of the YAC containing the viral genome in 7 out of 10 clones. kb, kilobases; pT7, T7 RNA polymerase promoter; An, poly (A) tail.

Extended Data Figure 1

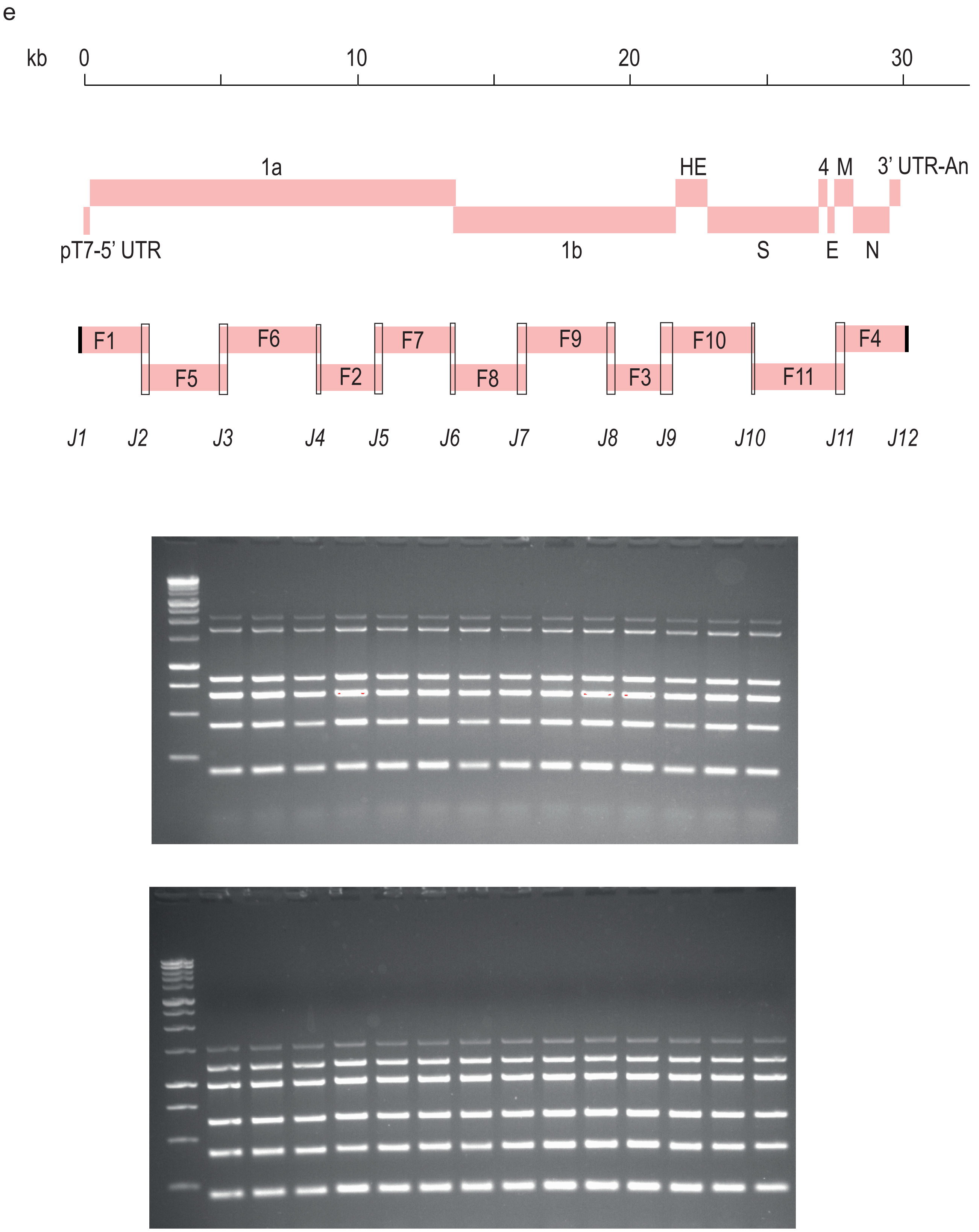


**e,** Schematic representation of the HCoV-HKU1 genome organisation (upper panel) and eleven viral subgenomic cDNA fragments (F1-11) used for TAR cloning. Viral open reading frames (ORFs) and sequence elements at the 5'- and 3'-untranslated regions (UTRs) are indicated. Primers used to generate the fragments are listed in Extended Data Table 1. J2-11 represent the junctions, i.e. overlapping regions, between the subgenomic cDNA fragments. J1 and J12 represent junctions with the TAR vector. Gel images show results from two multiplex PCRs that were designed to detect the presence of correctly recombined junctions J1-12, confirming the proper assembly of the YAC containing the viral genome in 14 out of 14 clones. kb, kilobases; pT7, T7 RNA polymerase promoter; An, poly (A) tail.

Extended Data Figure 1

f

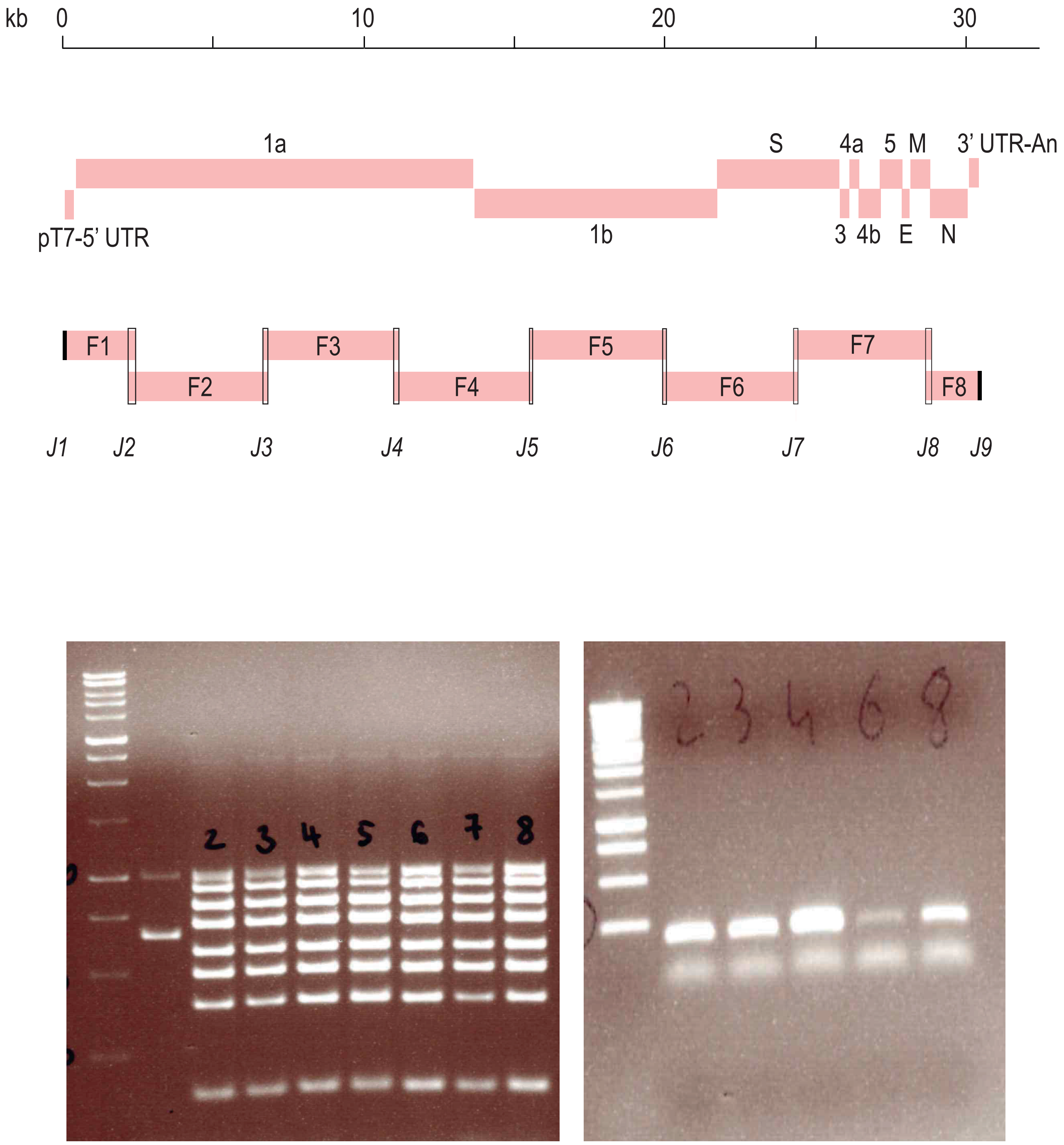


**f,** Schematic representation of the MERS-CoV-Riadh-1734-2015 genome organisation (upper panel) and eight viral subgenomic cDNA fragments (F1-8) used for TAR cloning. Viral open reading frames (ORFs) and sequence elements at the 5'- and 3'-untranslated regions (UTRs) are indicated. Primers used to generate the fragments are listed in Extended Data Table 1. J2-8 represent the junctions, i.e. overlapping regions, between the subgenomic cDNA fragments. J1 and J9 represent junctions with the TAR vector. Gel images show results from a multiplex PCR and a simplex PCR, both of which were designed to detect the presence of correctly recombined junctions J1-9, confirming the proper assembly of the YAC containing the viral genome in 5 out of 8 clones. kb, kilobases; pT7, T7 RNA polymerase promoter; An, poly (A) tail.

Extended Data Figure 1

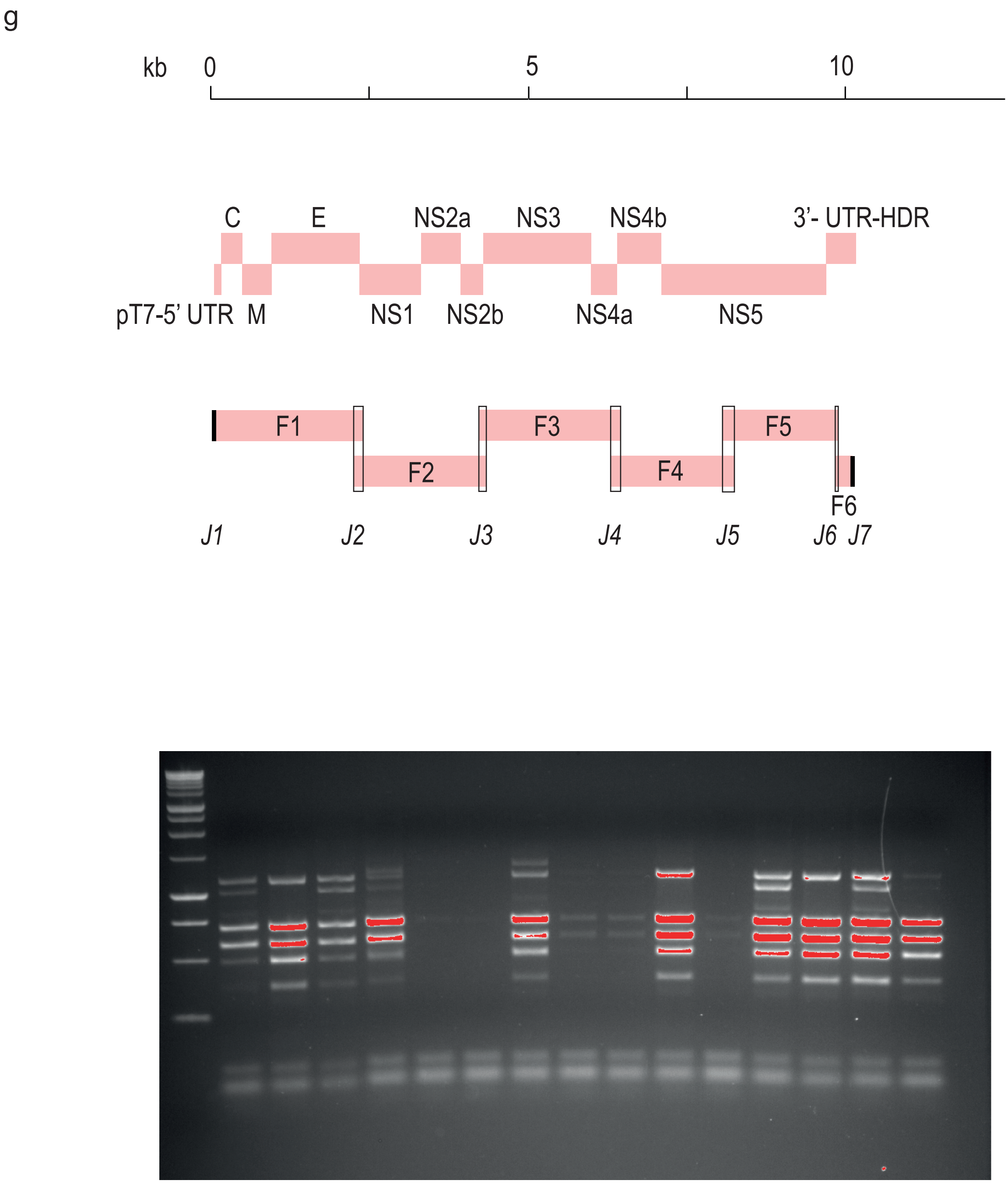


**g,** Schematic representation of the ZIKA virus genome organisation (upper panel) and six viral subgenomic cDNA fragments (F1-6) used for TAR cloning. Viral open reading frames (ORFs) and sequence elements at the 5'- and 3'-untranslated regions (UTRs) are indicated. Primers used to generate the fragments are listed in Extended Data Table 1. J2-5 represent the junctions, i.e. overlapping regions, between the subgenomic cDNA fragments. J1 and J6 represent junctions with the TAR vector. Gel image shows results from a multiplex PCR that was designed to detect the presence of correctly recombined junctions J1-6, confirming the proper assembly of the YAC containing the viral genome in 3 out of 15 clones. kb, kilobases; pT7, T7 RNA polymerase promoter; HDR, hepatitis delta virus ribozyme.

Extended Data Figure 1

h

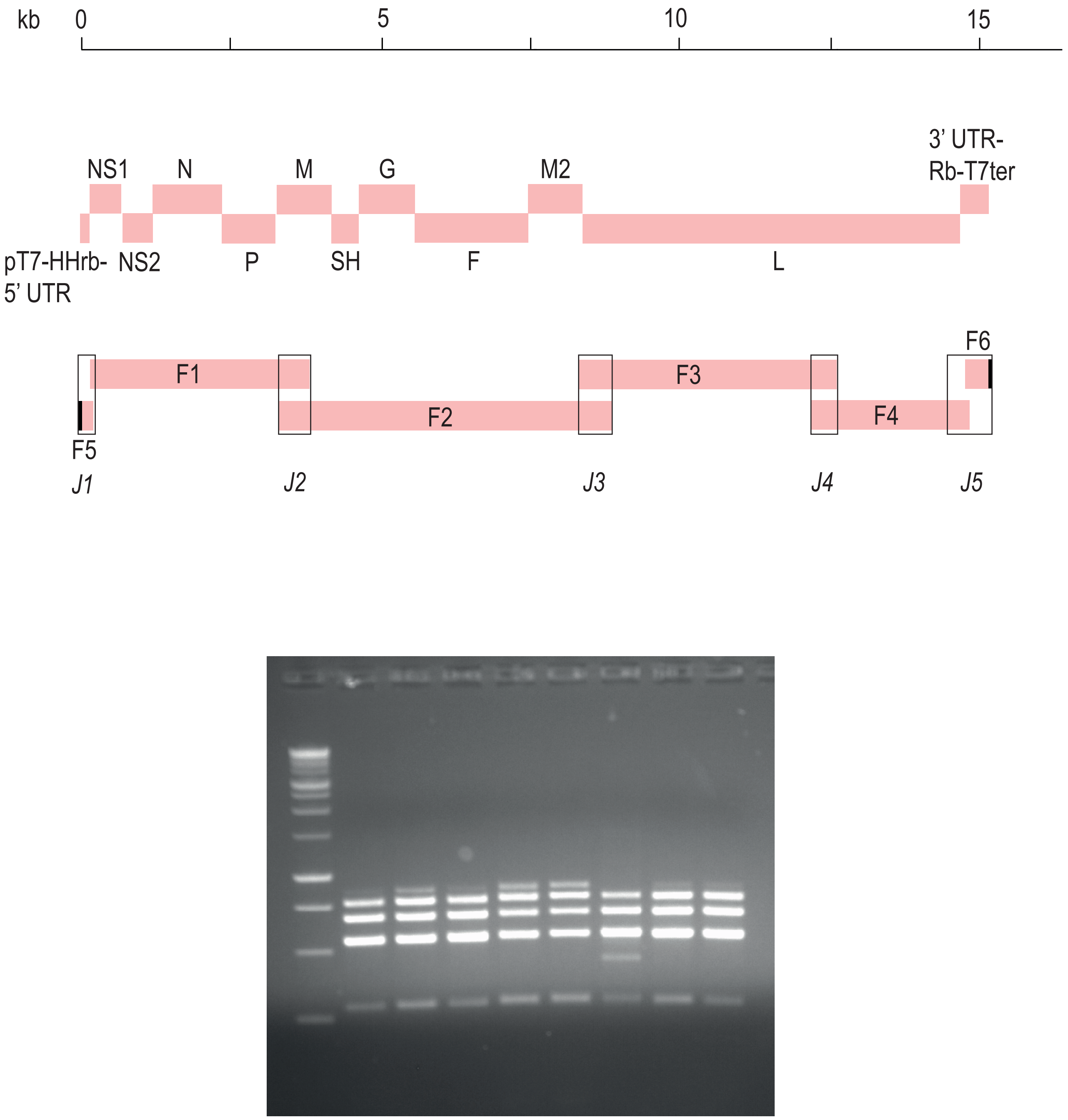


**h,** Schematic representation of the human RSV-B virus genome organisation (upper panel) and six viral subgenomic cDNA fragments (F1-6) used for TAR cloning. Viral open reading frames (ORFs) and sequence elements at the 5'- and 3'-untranslated regions (UTRs) are indicated. Primers used to generate the fragments are listed in Extended Data Table 1. J1-5 represent the junctions, i.e. overlapping regions, between the subgenomic cDNA fragments. J1 and J5 contain junctions with the TAR vector. Gel image shows results from a multiplex PCR that was designed to detect the presence of correctly recombined junctions J1-5, confirming the proper assembly of the YAC containing the viral genome in 7 out of 8 clones. kb, kilobases; pT7, T7 RNA polymerase promoter; HHrb, hammerhead ribozyme, Rb-T7ter, Ribozyme and T7 terminater.

### Extended Data Figure 1

i

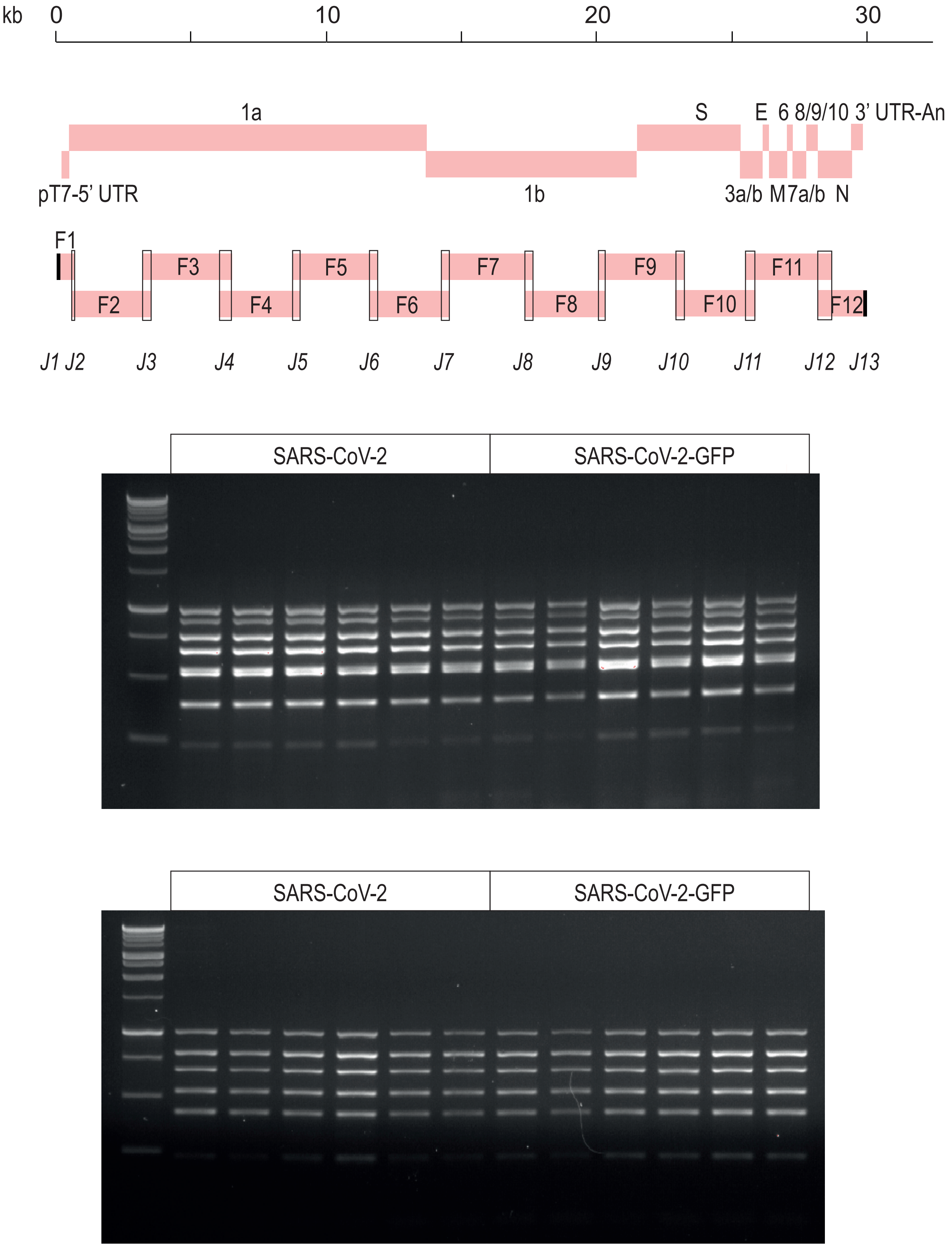


**i,** Schematic representation of the SARS-CoV-2 genome organisation (upper panel) and twelve viral subgenomic cDNA fragments (F1-12) used for TAR cloning. Viral open reading frames (ORFs) and sequence elements at the 5'- and 3'-untranslated regions (UTRs) are indicated. Primers used to generate the fragments are listed in Extended Data Table 1. J2-12 represent the junctions, i.e. overlapping regions, between the subgenomic cDNA fragments. J1 and J13 represent junctions with the TAR vector. Gel images show results from two multiplex PCRs that were designed to detect the presence of correctly recombined junctions J1-13, confirming the proper assembly of the YAC containing the viral genome in 6 out of 6 clones. kb, kilobases; pT7, T7 RNA polymerase promoter; An, poly (A) tail.

Extended Data Figure 1

j

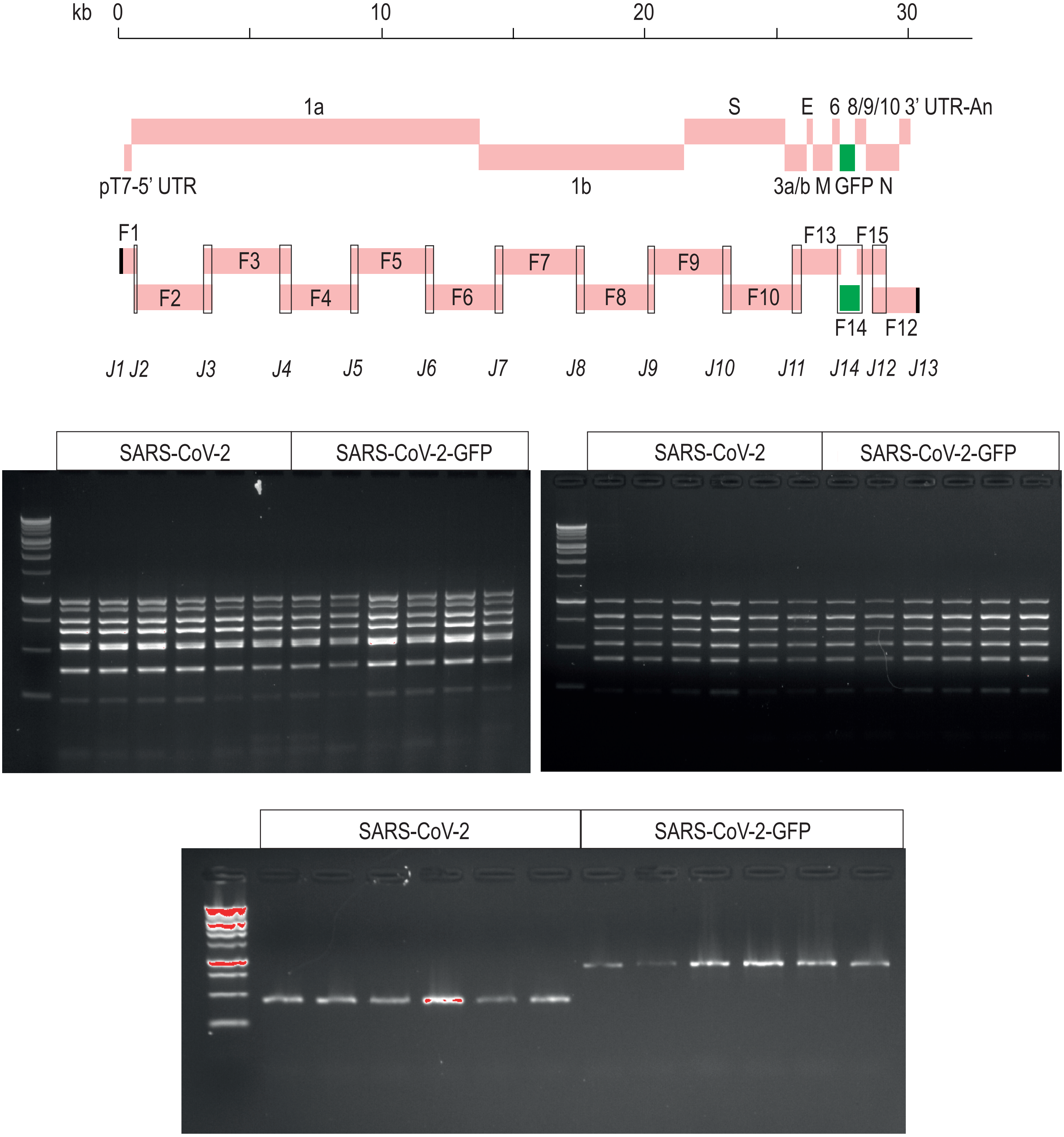


**j,** Schematic representation of the SARS-CoV-2 genome organisation (upper panel) and fourteen viral subgenomic cDNA fragments (F1-10 and F12-15) used for TAR cloning. Viral open reading frames (ORFs), ORF for GFP and sequence elements at the 5'- and 3'-untranslated regions (UTRs) are indicated. Primers used to generate the fragments are listed in Extended Data Table 1. J2-12 and J14 represent the junctions, i.e. overlapping regions, between the subgenomic cDNA fragments. J1 and J13 represent junctions with the TAR vector. Gel images show results from two multiplex PCRs and a simplex PCR, all of which were designed to detect the presence of correctly recombined junctions J1-13 and J14 respectively, confirming the proper assembly of the YAC containing the viral genome in 6 out of 6 clones. kb, kilobases; pT7, T7 RNA polymerase promoter; An, poly (A) tail.
