## Supplementary figures and images for "Rapid reconstruction of SARS-CoV-2 using a synthetic genomics platform"

### Extended Data Figure 2

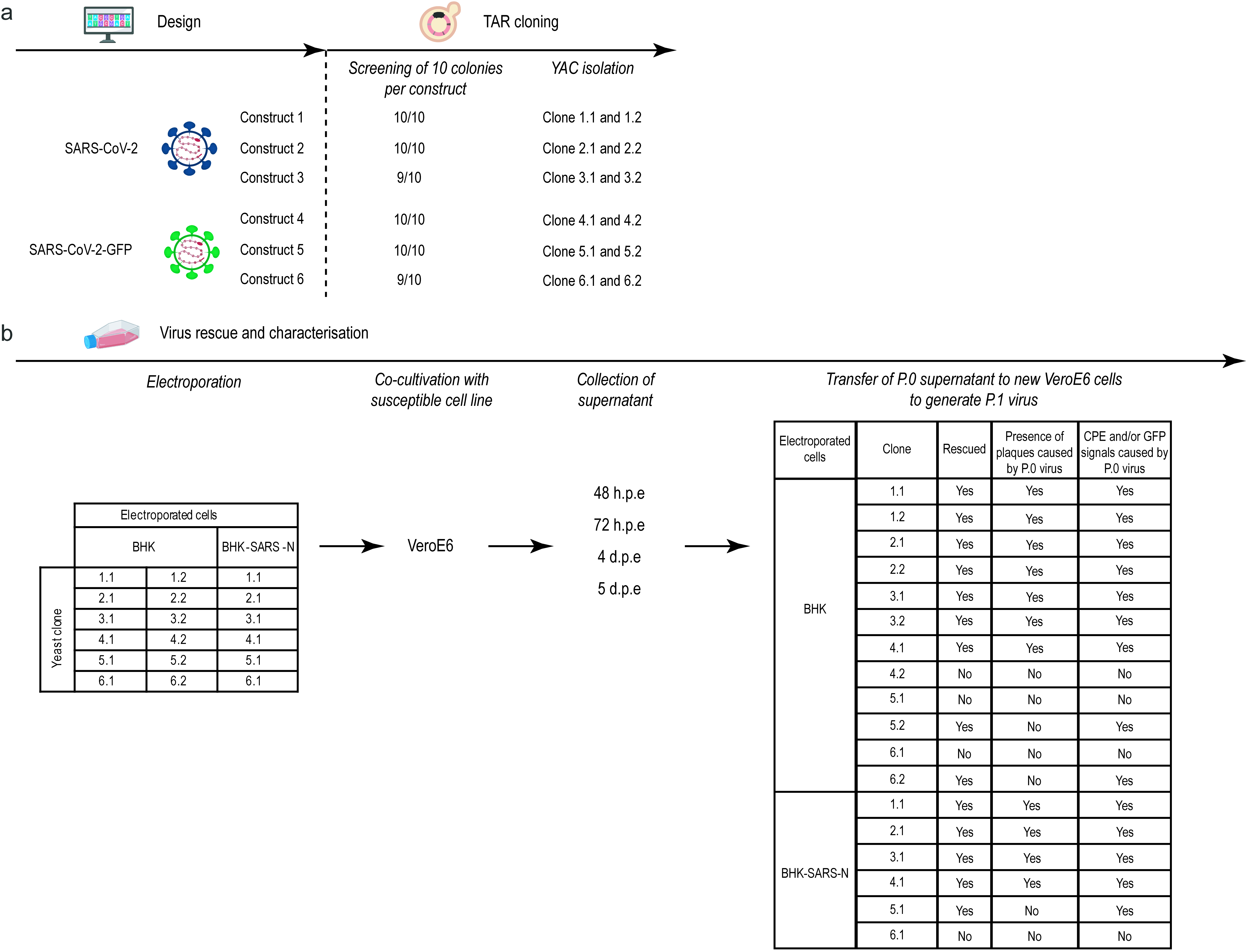
