## Extended Data Document 1 for "Rapid reconstruction of SARS-CoV-2 using a synthetic genomics platform"

SARS-CoV-2 synthetic fragments

Fragment 1.1

CCCG**ggcgaaagggggatgtgctgcaaggcgattaagttgggtaacgccagggttttcccagtcacgac catgcgtccatagtcccgttccg***TAATACGACTCACTATAG***ATATTAGGTTTATACCTTCCCAGGTAACAAACCAACCAACTTTCGATCTCTTGTAGATCTGTTCTCTAAACGAACTTTAAAATCTGTGTGGCTGTCACTCGGCTGCATGCTTAGTGCACTCACGCAGTATAATTAATAACTAATTACTGTCGTTGACAGGACACGAGTAACTCGTCTATCTTCTGCAGGCTGCTTACGGTTTCGTCCGTGTTGCAGCCGATCATCAGCACATCTAGGTTTCGTCCGGGTGTGACCGAAAGGTAAGATGGAGAGCCTTGTCCCTGGTTTCAACGAGAAAACACACGTCCAACTCAGTTTGCCTGTTTTACAGGTTCGCGACGTGCTCGTACGTGGCTTTGGAGACTCCGTGGAGGAGGTCTTATCAGAGGCACGTCAACATCTTAAAGATGGCACTTGTGGCTTAGTAGAAGTTGAAAAAGGCGTTTTGCCTCAACTTGAACAGCCCTATGTGTTCATCAAA**CCCGGG

Underline: *Sma*I

Lower case, bold: pCC1 and vaccinia virus sequences

Italic: T7 promoter

Bold: SARS-CoV-2, position 1 – 483. The viral 5'-UTR contains three modified nucleotides indicated in red.

Fragment 1.2

CCCG**ggcgaaagggggatgtgctgcaaggcgattaagttgggtaacgccagggttttcccagtcacgac catgcgtccatagtcccgttccg***TAATACGACTCACTATAG***TATTAGGTTTTTACCTACCCAGGAAAAGCCAACCAACCTCGATCTCTTGTAGATCTGTTCTCTAAACGAACTTTAAAATCTGTGTAGCTGTCGCTCGGCTGCATGCCTAGTGCACCTACGCAGTATAATTAATAACTAATTACTGTCGTTGACAGGACACGAGTAACTCGTCTATCTTCTGCAGGCTGCTTACGGTTTCGTCCGTGTTGCAGCCGATCATCAGCACATCTAGGTTTCGTCCGGGTGTGACCGAAAGGTAAGATGGAGAGCCTTGTCCCTGGTTTCAACGAGAAAACACACGTCCAACTCAGTTTGCCTGTTTTACAGGTTCGCGACGTGCTCGTACGTGGCTTTGGAGACTCCGTGGAGGAGGTCTTATCAGAGGCACGTCAACATCTTAAAGATGGCACTTGTGGCTTAGTAGAAGTTGAAAAAGGCGTTTTGCCTCAACTTGAACAGCCCTATGTGTTCATCAAA**CCCGGG

Underline: *Sma*I

Lower case, bold: pCC1 and vaccinia virus sequences

Italic: T7 promoter

Bold, underline: SARS-CoV, position 1 – 124

Bold: SARS-CoV-2, position 125 – 483

Fragment 1.3

CCCG**ggcgaaagggggatgtgctgcaaggcgattaagttgggtaacgccagggttttcccagtcacgac catgcgtccatagtcccgttccg** *TAATACGACTCACTATAG***ATTAAAGGTTTATACCTTCCCAGGTAACAAACCAACCAACTTTCGATCTCTTGTAGATCTGTTCTCTAAACGAACTTTAAAATCTGTGTGGCTGTCACTCGGCTGCATGCTTAGTGCACTCACGCAGTATAATTAATAACTAATTACTGTCGTTGACAGGACACGAGTAACTCGTCTATCTTCTGCAGGCTGCTTACGGTTTCGTCCGTGTTGCAGCCGATCATCAGCACATCTAGGTTTCGTCCGGGTGTGACCGAAAGGTAAGATGGAGAGCCTTGTCCCTGGTTTCAACGAGAAAACACACGTCCAACTCAGTTTGCCTGTTTTACAGGTTCGCGACGTGCTCGTACGTGGCTTTGGAGACTCCGTGGAGGAGGTCTTATCAGAGGCACGTCAACATCTTAAAGATGGCACTTGTGGCTTAGTAGAAGTTGAAAAAGGCGTTTTGCCTCAACTTGAACAGCCCTATGTGTTCATCAAA**CCCGGG

Underline: *Sma*I

Lower case, bold: pCC1 and vaccinia virus sequences

Italic: T7 promoter

Bold: SARS-CoV-2, position 1 – 483.

Fragment 2

CCCGG**GGTCTTATCAGAGGCACGTCAACATCTTAAAGATGGCACTTGTGGCTTAGTAGAAGTTGAAAAAGGCGTTTTGCCTCAACTTGAACAGCCCTATGTGTTCATCAAACGTTCGGATGCTCGAACTGCACCTCATGGTCATGTTATGGTTGAGCTGGTAGCAGAACTCGAAGGCATTCAGTACGGTCGTAGTGGTGAGACACTTGGTGTCCTTGTCCCTCATGTGGGCGAAATACCAGTGGCTTACCGCAAGGTTCTTCTTCGTAAGAACGGTAATAAAGGAGCTGGTGGCCATAGTTACGGCGCCGATCTAAAGTCATTTGACTTAGGCGACGAGCTTGGCACTGATCCTTATGAAGATTTTCAAGAAAACTGGAACACTAAACATAGCAGTGGTGTTACCCGTGAACTCATGCGTGAGCTTAACGGAGGGGCATACACTCGCTATGTCGATAACAACTTCTGTGGCCCTGATGGCTACCCTCTTGAGTGCATTAAAGACCTTCTAGCACGTGCTGGTAAAGCTTCATGCACTTTGTCCGAACAACTGGACTTTATTGACACTAAGAGGGGTGTATACTGCTGCCGTGAACATGAGCATGAAATTGCTTGGTACACGGAACGTTCTGAAAAGAGCTATGAATTGCAGACACCTTTTGAAATTAAATTGGCAAAGAAATTTGACACCTTCAATGGGGAATGTCCAAATTTTGTATTTCCCTTAAATTCCATAATCAAGACTATTCAACCAAGGGTTGAAAAGAAAAAGCTTGATGGCTTTATGGGTAGAATTCGATCTGTCTATCCAGTTGCGTCACCAAATGAATGCAACCAAATGTGCCTTTCAACTCTCATGAAGTGTGATCATTGTGGTGAAACTTCATGGCAGACGGGCGATTTTGTTAAAGCCACTTGCGAATTTTGTGGCACTGAGAATTTGACTAAAGAAGGTGCCACTACTTGTGGTTACTTACCCCAAAATGCTGTTGTTAAAATTTATTGTCCAGCATGTCACAATTCAGAAGTAGGACCTGAGCATAGTCTTGCCGAATACCATAATGAATCTGGCTTGAAAACCATTCTTCGTAAGGGTGGTCGCACTATTGCCTTTGGAGGCTGTGTGTTCTCTTATGTTGGTTGCCATAACAAGTGTGCCTATTGGGTTCCACGTGCTAGCGCTAACATAGGTTGTAACCATACAGGTGTTGTTGGAGAAGGTTCCGAAGGTCTTAATGACAACCTTCTTGAAATACTCCAAAAAGAGAAAGTCAACATCAATATTGTTGGTGACTTTAAACTTAATGAAGAGATCGCCATTATTTTGGCATCTTTTTCTGCTTCCACAAGTGCTTTTGTGGAAACTGTGAAAGGTTTGGATTATAAAGCATTCAAACAAATTGTTGAATCCTGTGGTAATTTTAAAGTTACAAAAGGAAAAGCTAAAAAAGGTGCCTGGAATATTGGTGAACAGAAATCAATACTGAGTCCTCTTTATGCATTTGCATCAGAGGCTGCTCGTGTTGTACGATCAATTTTCTCCCGCACTCTTGAAACTGCTCAAAATTCTGTGCGTGTTTTACAGAAGGCCGCTATAACAATACTAGATGGAATTTCACAGTATTCACTGAGACTCATTGATGCTATGATGTTCACATCTGATTTGGCTACTAACAATCTAGTTGTAATGGCCTACATTACAGGTGGTGTTGTTCAGTTGACTTCGCAGTGGCTAACTAACATCTTTGGCACTGTTTATGAAAAACTCAAACCCGTCCTTGATTGGCTTGAAGAGAAGTTTAAGGAAGGTGTAGAGTTTCTTAGAGACGGTTGGGAAATTGTTAAATTTATCTCAACCTGTGCTTGTGAAATTGTCGGTGGACAAATTGTCACCTGTGCAAAGGAAATTAAGGAGAGTGTTCAGACATTCTTTAAGCTTGTAAATAAATTTTTGGCTTTGTGTGCTGACTCTATCATTATTGGTGGAGCTAAACTTAAAGCCTTGAATTTAGGTGAAACATTTGTCACGCACTCAAAGGGATTGTACAGAAAGTGTGTTAAATCCAGAGAAGAAACTGGCCTACTCATGCCTCTAAAAGCCCCAAAAGAAATTATCTTCTTAGAGGGAGAAACACTTCCCACAGAAGTGTTAACAGAGGAAGTTGTCTTGAAAACTGGTGATTTACAACCATTAGAACAACCTACTAGTGAAGCTGTTGAAGCTCCATTGGTTGGTACACCAGTTTGTATTAACGGGCTTATGTTGCTCGAAATCAAAGACACAGAAAAGTACTGTGCCCTTGCACCTAATATGATGGTAACAAACAATACCTTCACACTCAAAGGCGGTGCACCAACAAAGGTTACTTTTGGTGATGACACTGTGATAGAAGTGCAAGGTTACAAGAGTGTGAATATCACTTTTGAACTTGATGAAAGGATTGATAAAGTACTTAATGAGAAGTGCTCTGCCTATACAGTTGAACTCGGTACAGAAGTAAATGAGTTCGCCTGTGTTGTGGCAGATGCTGTCATAAAAACTTTGCAACCAGTATCTGAATTACTTACACCACTGGGCATTGATTTAGATGAGTGGAGTATGGCTACATACTACTTATTTGATGAGTCTGGTGAGTTTAAATTGGCTTCACATATGTATTGTTCTTTCTACCCTCCAGATGAGGATGAAGAAGAAGGTGATTGTGAAGAAGAAGAGTTTGAGCCATCAACTCAATATGAGTATGGTACTGAAGATGATTACCAAGGTAAACCTTTGGAATTTGGTGCCACTTCTGCTGCTCTTCAACCTGAAGAAGAGCAAGAAGAAGATTGGTTAGATGATGATAGTCAACAAACTGTTGGTCAACAAGACGGCAGTGAGGACAATCAGACAACTACTATTCAAACAATTGTTGAGGTTCAACCTCAATTAGAGATGGAACTTACACCAGTTGTT**CCCGGG

Underline: *Sma*I

Bold: SARS-CoV-2, position 377 – 3325

Fragment 3

GATATC**TGGCTTCACATATGTATTGTTCTTTCTACCCTCCAGATGAGGATGAAGAAGAAGGTGATTGTGAAGAAGAAGAGTTTGAGCCATCAACTCAATATGAGTATGGTACTGAAGATGATTACCAAGGTAAACCTTTGGAATTTGGTGCCACTTCTGCTGCTCTTCAACCTGAAGAAGAGCAAGAAGAAGATTGGTTAGATGATGATAGTCAACAAACTGTTGGTCAACAAGACGGCAGTGAGGACAATCAGACAACTACTATTCAAACAATTGTTGAGGTTCAACCTCAATTAGAGATGGAACTTACACCAGTTGTTCAGACTATTGAAGTGAATAGTTTTAGTGGTTATTTAAAACTTACTGACAATGTATACATTAAAAATGCAGACATTGTGGAAGAAGCTAAAAAGGTAAAACCAACAGTGGTTGTTAATGCAGCCAATGTTTACCTTAAACATGGAGGAGGTGTTGCAGGAGCCTTAAATAAGGCTACTAACAATGCCATGCAAGTTGAATCTGATGATTACATAGCTACTAATGGACCACTTAAAGTGGGTGGTAGTTGTGTTTTAAGCGGACACAATCTTGCTAAACACTGTCTTCATGTTGTCGGCCCAAATGTTAACAAAGGTGAAGACATTCAACTTCTTAAGAGTGCTTATGAAAATTTTAATCAGCACGAAGTTCTACTTGCACCATTATTATCAGCTGGTATTTTTGGTGCTGACCCTATACATTCTTTAAGAGTTTGTGTAGATACTGTTCGCACAAATGTCTACTTAGCTGTCTTTGATAAAAATCTCTATGACAAACTTGTTTCAAGCTTTTTGGAAATGAAGAGTGAAAAGCAAGTTGAACAAAAGATCGCTGAGATTCCTAAAGAGGAAGTTAAGCCATTTATAACTGAAAGTAAACCTTCAGTTGAACAGAGAAAACAAGATGATAAGAAAATCAAAGCTTGTGTTGAAGAAGTTACAACAACTCTGGAAGAAACTAAGTTCCTCACAGAAAACTTGTTACTTTATATTGACATTAATGGCAATCTTCATCCAGATTCTGCCACTCTTGTTAGTGACATTGACATCACTTTCTTAAAGAAAGATGCTCCATATATAGTGGGTGATGTTGTTCAAGAGGGTGTTTTAACTGCTGTGGTTATACCTACTAAAAAGGCTGGTGGCACTACTGAAATGCTAGCGAAAGCTTTGAGAAAAGTGCCAACAGACAATTATATAACCACTTACCCGGGTCAGGGTTTAAATGGTTACACTGTAGAGGAGGCAAAGACAGTGCTTAAAAAGTGTAAAAGTGCCTTTTACATTCTACCATCTATTATCTCTAATGAGAAGCAAGAAATTCTTGGAACTGTTTCTTGGAATTTGCGAGAAATGCTTGCACATGCAGAAGAAACACGCAAATTAATGCCTGTCTGTGTGGAAACTAAAGCCATAGTTTCAACTATACAGCGTAAATATAAGGGTATTAAAATACAAGAGGGTGTGGTTGATTATGGTGCTAGATTTTACTTTTACACCAGTAAAACAACTGTAGCGTCACTTATCAACACACTTAACGATCTAAATGAAACTCTTGTTACAATGCCACTTGGCTATGTAACACATGGCTTAAATTTGGAAGAAGCTGCTCGGTATATGAGATCTCTCAAAGTGCCAGCTACAGTTTCTGTTTCTTCACCTGATGCTGTTACAGCGTATAATGGTTATCTTACTTCTTCTTCTAAAACACCTGAAGAACATTTTATTGAAACCATCTCACTTGCTGGTTCCTATAAAGATTGGTCCTATTCTGGACAATCTACACAACTAGGTATAGAATTTCTTAAGAGAGGTGATAAAAGTGTATATTACACTAGTAATCCTACCACATTCCACCTAGATGGTGAAGTTATCACCTTTGACAATCTTAAGACACTTCTTTCTTTGAGAGAAGTGAGGACTATTAAGGTGTTTACAACAGTAGACAACATTAACCTCCACACGCAAGTTGTGGACATGTCAATGACATATGGACAACAGTTTGGTCCAACTTATTTGGATGGAGCTGATGTTACTAAAATAAAACCTCATAATTCACATGAAGGTAAAACATTTTATGTTTTACCTAATGATGACACTCTACGTGTTGAGGCTTTTGAGTACTACCACACAACTGATCCTAGTTTTCTGGGTAGGTACATGTCAGCATTAAATCACACTAAAAAGTGGAAATACCCACAAGTTAATGGTTTAACTTCTATTAAATGGGCAGATAACAACTGTTATCTTGCCACTGCATTGTTAACACTCCAACAAATAGAGTTGAAGTTTAATCCACCTGCTCTACAAGATGCTTATTACAGAGCAAGGGCTGGTGAAGCTGCTAACTTTTGTGCACTTATCTTAGCCTACTGTAATAAGACAGTAGGTGAGTTAGGTGATGTTAGAGAAACAATGAGTTACTTGTTTCAACATGCCAATTTAGATTCTTGCAAAAGAGTCTTGAACGTGGTGTGTAAAACTTGTGGACAACAGCAGACAACCCTTAAGGGTGTAGAAGCTGTTATGTACATGGGCACACTTTCTTATGAACAATTTAAGAAAGGTGTTCAGATACCTTGTACGTGTGGTAAACAAGCTACAAAATATCTAGTACAACAGGAGTCACCTTTTGTTATGATGTCAGCACCACCTGCTCAGTATGAACTTAAGCATGGTACATTTACTTGTGCTAGTGAGTACACTGGTAATTACCAGTGTGGTCACTATAAACATATAACTTCTAAAGAAACTTTGTATTGCATAGACGGTGCTTTACTTACAAAGTCCTCAGAATACAAAGGTCCTATTACGGATGTTTTCTACAAAGAAAACAGTTACACAACAACCATAAAACCAGTTACTTATAAATTGGATGGTGTTGTTTGTACAGAAATTGACCCTAAGTTGGACAATTATTATAAGAAAGACAATTCTTATTTCACAGAGCAACCAATTGATCTTGTACCAAACCAACCATATCCAAACGCAAGCTTCGATAATTTTAAGTTTGTATGTGATAATATCAAATTTGCTGATGATTTAAACCAGTTAACTGGTTATAAGAAACCTGCTTCAAGAGAGCTTAAAGTTACATTTTTCCCTGACTTAAATGGTGATGTGGTGGCTATTGATTATAAACACTACACACCCTCTTTTAAGAAAGGAGCTAAATTGTTACATAAACCTATTGTTTGGCATGTTAACAATGCAACTAATAAAGCCACGTATAAACCAAATACCTGGTGTATACGTTGTCTTTGGAGCACAAA**GATATC

Underline: *Eco*RV

Bold: SARS-CoV-2, position 3012 – 6315

Fragment 4

GATATC**TTGTACCAAACCAACCATATCCAAACGCAAGCTTCGATAATTTTAAGTTTGTATGTGATAATATCAAATTTGCTGATGATTTAAACCAGTTAACTGGTTATAAGAAACCTGCTTCAAGAGAGCTTAAAGTTACATTTTTCCCTGACTTAAATGGTGATGTGGTGGCTATTGATTATAAACACTACACACCCTCTTTTAAGAAAGGAGCTAAATTGTTACATAAACCTATTGTTTGGCATGTTAACAATGCAACTAATAAAGCCACGTATAAACCAAATACCTGGTGTATACGTTGTCTTTGGAGCACAAAACCAGTTGAAACATCAAATTCGTTTGATGTACTGAAGTCAGAGGACGCGCAGGGAATGGATAATCTTGCCTGCGAAGATCTAAAACCAGTCTCTGAAGAAGTAGTGGAAAATCCTACCATACAGAAAGACGTTCTTGAGTGTAATGTGAAAACTACCGAAGTTGTAGGAGACATTATACTTAAACCAGCAAATAATAGTTTAAAAATTACAGAAGAGGTTGGCCACACAGATCTAATGGCTGCTTATGTAGACAATTCTAGTCTTACTATTAAGAAACCTAATGAATTATCTAGAGTATTAGGTTTGAAAACCCTTGCTACTCATGGTTTAGCTGCTGTTAATAGTGTCCCTTGGGATACTATAGCTAATTATGCTAAGCCTTTTCTTAACAAAGTTGTTAGTACAACTACTAACATAGTTACACGGTGTTTAAACCGTGTTTGTACTAATTATATGCCTTATTTCTTTACTTTATTGCTACAATTGTGTACTTTTACTAGAAGTACAAATTCTAGAATTAAAGCATCTATGCCGACTACTATAGCAAAGAATACTGTTAAGAGTGTCGGTAAATTTTGTCTAGAGGCTTCATTTAATTATTTGAAGTCACCTAATTTTTCTAAACTGATAAATATTATAATTTGGTTTTTACTATTAAGTGTTTGCCTAGGTTCTTTAATCTACTCAACCGCTGCTTTAGGTGTTTTAATGTCTAATTTAGGCATGCCTTCTTACTGTACTGGTTACAGAGAAGGCTATTTGAACTCTACTAATGTCACTATTGCAACCTACTGTACTGGTTCTATACCTTGTAGTGTTTGTCTTAGTGGTTTAGATTCTTTAGACACCTATCCTTCTTTAGAAACTATACAAATTACCATTTCATCTTTTAAATGGGATTTAACTGCTTTTGGCTTAGTTGCAGAGTGGTTTTTGGCATATATTCTTTTCACTAGGTTTTTCTATGTACTTGGATTGGCTGCAATCATGCAATTGTTTTTCAGCTATTTTGCAGTACATTTTATTAGTAATTCTTGGCTTATGTGGTTAATAATTAATCTTGTACAAATGGCCCCGATTTCAGCTATGGTTAGAATGTACATCTTCTTTGCATCATTTTATTATGTATGGAAAAGTTATGTGCATGTTGTAGACGGTTGTAATTCATCAACTTGTATGATGTGTTACAAACGTAATAGAGCAACAAGAGTCGAATGTACAACTATTGTTAATGGTGTTAGAAGGTCCTTTTATGTCTATGCTAATGGAGGTAAAGGCTTTTGCAAACTACACAATTGGAATTGTGTTAATTGTGATACATTCTGTGCTGGTAGTACATTTATTAGTGATGAAGTTGCGAGAGACTTGTCACTACAGTTTAAAAGACCAATAAATCCTACTGACCAGTCTTCTTACATCGTTGATAGTGTTACAGTGAAGAATGGTTCCATCCATCTTTACTTTGATAAAGCTGGTCAAAAGACTTATGAAAGACATTCTCTCTCTCATTTTGTTAACTTAGACAACCTGAGAGCTAATAACACTAAAGGTTCATTGCCTATTAATGTTATAGTTTTTGATGGTAAATCAAAATGTGAAGAATCATCTGCAAAATCAGCGTCTGTTTACTACAGTCAGCTTATGTGTCAACCTATACTGTTACTAGATCAGGCATTAGTGTCTGATGTTGGTGATAGTGCGGAAGTTGCAGTTAAAATGTTTGATGCTTACGTTAATACGTTTTCATCAACTTTTAACGTACCAATGGAAAAACTCAAAACACTAGTTGCAACTGCAGAAGCTGAACTTGCAAAGAATGTGTCCTTAGACAATGTCTTATCTACTTTTATTTCAGCAGCTCGGCAAGGGTTTGTTGATTCAGATGTAGAAACTAAAGATGTTGTTGAATGTCTTAAATTGTCACATCAATCTGACATAGAAGTTACTGGCGATAGTTGTAATAACTATATGCTCACCTATAACAAAGTTGAAAACATGACACCCCGTGACCTTGGTGCTTGTATTGACTGTAGTGCGCGTCATATTAATGCGCAGGTAGCAAAAAGTCACAACATTGCTTTGATATGGAACGTTAAAGATTTCATGTCATTGTCTGAACAACTACGAAAACAAATACGTAGTGCTGCTAAAAAGAATAACTTACCTTTTAAGTTGACATGTGCAACTACTAGACAAGTTGTTAATGTTGTAACAACAAAGATAGCACTTAAGGGTGGTAAAATTGTTAATAATTGGTTGAAGCAGTTAATTAAAGTTACACTTGTGTTCCTTTTTGTTGCTGCTATTTTCTATTTAATAACACCTGTTCATGTCATGTCTAAACATACTGACTTTTCAAGTGAAATCATAGGATACAAGGCTATTGATGGTGGTGTCACTCGTGACATAGCATCTACAGATACTTGTTTTGCTAACAAACATGCTGATTTTGACACATGGTTTAGCCAGCGTGGTGGTAGTTATACTAATGACAAAGCTTGCCCATTGATTGCTGCAGTCATAACAAGAGAAGTGGGTTTTGTCGTGCCTGGTTTGCCTGGCACGATATTACGCACAACTAATGGTGACTTTTTGCATTTCTTACCTAGAGTTTTTAGTGCAGTTGGTAACATCTGTTACACACCATCAAAACTTATAGAGTACACTGACTTTGCAA*C****ATATG*

Underline: *Eco*RV

Underline, italic: *Nde*I

Bold: SARS-CoV-2, position 6003 – 8994

Fragment 5

CCCGGG**TGACATAGCATCTACAGATACTTGTTTTGCTAACAAACATGCTGATTTTGACACATGGTTTAGCCAGCGTGGTGGTAGTTATACTAATGACAAAGCTTGCCCATTGATTGCTGCAGTCATAACAAGAGAAGTGGGTTTTGTCGTGCCTGGTTTGCCTGGCACGATATTACGCACAACTAATGGTGACTTTTTGCATTTCTTACCTAGAGTTTTTAGTGCAGTTGGTAACATCTGTTACACACCATCAAAACTTATAGAGTACACTGACTTTGCAACATCAGCTTGTGTTTTGGCTGCTGAATGTACAATTTTTAAAGATGCTTCTGGTAAGCCAGTACCATATTGTTATGATACCAATGTACTAGAAGGTTCTGTTGCTTATGAAAGTTTACGCCCTGACACACGTTATGTGCTCATGGATGGCTCTATTATTCAATTTCCTAACACCTACCTTGAAGGTTCTGTTAGAGTGGTAACAACTTTTGATTCTGAGTACTGTAGGCACGGCACTTGTGAAAGATCAGAAGCTGGTGTTTGTGTATCTACTAGTGGTAGATGGGTACTTAACAATGATTATTACAGATCTTTACCAGGAGTTTTCTGTGGTGTAGATGCTGTAAATTTACTTACTAATATGTTTACACCACTAATTCAACCTATTGGTGCTTTGGACATATCAGCATCTATAGTAGCTGGTGGTATTGTAGCTATCGTAGTAACATGCCTTGCCTACTATTTTATGAGGTTTAGAAGAGCTTTTGGTGAATACAGTCATGTAGTTGCCTTTAATACTTTACTATTCCTTATGTCATTCACTGTACTCTGTTTAACACCAGTTTACTCATTCTTACCTGGTGTTTATTCTGTTATTTACTTGTACTTGACATTTTATCTTACTAATGATGTTTCTTTTTTAGCACATATTCAGTGGATGGTTATGTTCACACCTTTAGTACCTTTCTGGATAACAATTGCTTATATCATTTGTATTTCCACAAAGCATTTCTATTGGTTCTTTAGTAATTACCTAAAGAGACGTGTAGTCTTTAATGGTGTTTCCTTTAGTACTTTTGAAGAAGCTGCGCTGTGCACCTTTTTGTTAAATAAAGAAATGTATCTAAAGTTGCGTAGTGATGTGCTATTACCTCTTACGCAATATAATAGATACTTAGCTCTTTATAATAAGTACAAGTATTTTAGTGGAGCAATGGATACAACTAGCTACAGAGAAGCTGCTTGTTGTCATCTCGCAAAGGCTCTCAATGACTTCAGTAACTCAGGTTCTGATGTTCTTTACCAACCACCACAAACCTCTATCACCTCAGCTGTTTTGCAGAGTGGTTTTAGAAAAATGGCATTCCCATCTGGTAAAGTTGAGGGTTGTATGGTACAAGTAACTTGTGGTACAACTACACTTAACGGTCTTTGGCTTGATGACGTAGTTTACTGTCCAAGACATGTGATCTGCACCTCTGAAGACATGCTTAACCCTAATTATGAAGATTTACTCATTCGTAAGTCTAATCATAATTTCTTGGTACAGGCTGGTAATGTTCAACTCAGGGTTATTGGACATTCTATGCAAAATTGTGTACTTAAGCTTAAGGTTGATACAGCCAATCCTAAGACACCTAAGTATAAGTTTGTTCGCATTCAACCAGGACAGACTTTTTCAGTGTTAGCTTGTTACAATGGTTCACCATCTGGTGTTTACCAATGTGCTATGAGGCCCAATTTCACTATTAAGGGTTCATTCCTTAATGGTTCATGTGGTAGTGTTGGTTTTAACATAGATTATGACTGTGTCTCTTTTTGTTACATGCACCATATGGAATTACCAACTGGAGTTCATGCTGGCACAGACTTAGAAGGTAACTTTTATGGACCTTTTGTTGACAGGCAAACAGCACAAGCAGCTGGTACGGACACAACTATTACAGTTAATGTTTTAGCTTGGTTGTACGCTGCTGTTATAAATGGAGACAGGTGGTTTCTCAATCGATTTACCACAACTCTTAATGACTTTAACCTTGTGGCTATGAAGTACAATTATGAACCTCTAACACAAGACCATGTTGACATACTAGGACCTCTTTCTGCTCAAACTGGAATTGCCGTTTTAGATATGTGTGCTTCATTAAAAGAATTACTGCAAAATGGTATGAATGGACGTACCATATTGGGTAGTGCTTTATTAGAAGATGAATTTACACCTTTTGATGTTGTTAGACAATGCTCAGGTGTTACTTTCCAAAGTGCAGTGAAAAGAACAATCAAGGGTACACACCACTGGTTGTTACTCACAATTTTGACTTCACTTTTAGTTTTAGTCCAGAGTACTCAATGGTCTTTGTTCTTTTTTTTGTATGAAAATGCCTTTTTACCTTTTGCTATGGGTATTATTGCTATGTCTGCTTTTGCAATGATGTTTGTCAAACATAAGCATGCATTTCTCTGTTTGTTTTTGTTACCTTCTCTTGCCACTGTAGCTTATTTTAATATGGTCTATATGCCTGCTAGTTGGGTGATGCGTATTATGACATGGTTGGATATGGTTGATACTAGTTTGTCTGGTTTTAAGCTAAAAGACTGTGTTATGTATGCATCAGCTGTAGTGTTACTAATCCTTATGACAGCAAGAACTGTGTATGATGATGGTGCTAGGAGAGTGTGGACACTTATGAATGTCTTGACACTCGTTTATAAAGTTTATTATGGTAATGCTTTAGATCAAGCCATTTCCATGTGGGCTCTTATAATCTCTGTTACTTCTAACTACTCAGGTGTAGTTACAACTGTCATGTTTTTGGCCAGAGGTATTGTTTTTATGTGTGTTGAGTATTGCCCTATTTTCTTCATAACTGGTAATACACTTCAGTGTATAATGCTAGTTTATTGTTTCTTAGGCTATTTTTGTACTTGTTACTTTGGCCTCTTTTGTTTACTCAACCGCTACTTTAGACTGACTCTTGGTGTTTATGATTACTTAGTTTCTACACAGGAGTTTAGATATATGAATTCACAGGGACTACTCCCACCCAAGAATAGCATAGATGCCTTCAAACTCAACATTAAATTGTTGGGTGTTGGTGGCAAACCTTGTATCAAAGTAGCCACTGTACAGTCTAAAATGTCAGATGTAAAGTGCACATCAGTAGTCTTACTCTCAGTTTTGCAACAACTCAGAGTAGAATCATCATCTAAATTGTGGGCTCAATGTGTCCAGTTACACAATGACATTCTCTTAG**CCCGGG

Underline: *Sma*I

Bold: SARS-CoV-2, position 8718 – 11966

Fragment 6

CCCGGG**TCAACCGCTACTTTAGACTGACTCTTGGTGTTTATGATTACTTAGTTTCTACACAGGAGTTTAGATATATGAATTCACAGGGACTACTCCCACCCAAGAATAGCATAGATGCCTTCAAACTCAACATTAAATTGTTGGGTGTTGGTGGCAAACCTTGTATCAAAGTAGCCACTGTACAGTCTAAAATGTCAGATGTAAAGTGCACATCAGTAGTCTTACTCTCAGTTTTGCAACAACTCAGAGTAGAATCATCATCTAAATTGTGGGCTCAATGTGTCCAGTTACACAATGACATTCTCTTAGCTAAAGATACTACTGAAGCCTTTGAAAAAATGGTTTCACTACTTTCTGTTTTGCTTTCCATGCAGGGTGCTGTAGACATAAACAAGCTTTGTGAAGAAATGCTGGACAACAGGGCAACCTTACAAGCTATAGCCTCAGAGTTTAGTTCCCTTCCATCATATGCAGCTTTTGCTACTGCTCAAGAAGCTTATGAGCAGGCTGTTGCTAATGGTGATTCTGAAGTTGTTCTTAAAAAGTTGAAGAAGTCTTTGAATGTGGCTAAATCTGAATTTGACCGTGATGCAGCCATGCAACGTAAGTTGGAAAAGATGGCTGATCAAGCTATGACCCAAATGTATAAACAGGCTAGATCTGAGGACAAGAGGGCAAAAGTTACTAGTGCTATGCAGACAATGCTTTTCACTATGCTTAGAAAGTTGGATAATGATGCACTCAACAACATTATCAACAATGCAAGAGATGGTTGTGTTCCCTTGAACATAATACCTCTTACAACAGCAGCCAAACTAATGGTTGTCATACCAGACTATAACACATATAAAAATACGTGTGATGGTACAACATTTACTTATGCATCAGCATTGTGGGAAATCCAACAGGTTGTAGATGCAGATAGTAAAATTGTTCAACTTAGTGAAATTAGTATGGACAATTCACCTAATTTAGCATGGCCTCTTATTGTAACAGCTTTAAGGGCCAATTCTGCTGTCAAATTACAGAATAATGAGCTTAGTCCTGTTGCACTACGACAGATGTCTTGTGCTGCCGGTACTACACAAACTGCTTGCACTGATGACAATGCGTTAGCTTACTACAACACAACAAAGGGAGGTAGGTTTGTACTTGCACTGTTATCCGATTTACAGGATTTGAAATGGGCTAGATTCCCTAAGAGTGATGGAACTGGTACTATCTATACAGAACTGGAACCACCTTGTAGGTTTGTTACAGACACACCTAAAGGTCCTAAAGTGAAGTATTTATACTTTATTAAAGGATTAAACAACCTAAATAGAGGTATGGTACTTGGTAGTTTAGCTGCCACAGTACGTCTACAAGCTGGTAATGCAACAGAAGTGCCTGCCAATTCAACTGTATTATCTTTCTGTGCTTTTGCTGTAGATGCTGCTAAAGCTTACAAAGATTATCTAGCTAGTGGGGGACAACCAATCACTAATTGTGTTAAGATGTTGTGTACACACACTGGTACTGGTCAGGCAATAACAGTTACACCGGAAGCCAATATGGATCAAGAATCCTTTGGTGGTGCATCGTGTTGTCTGTACTGCCGTTGCCACATAGATCATCCAAATCCTAAAGGATTTTGTGACTTAAAAGGTAAGTATGTACAAATACCTACAACTTGTGCTAATGACCCTGTGGGTTTTACACTTAAAAACACAGTCTGTACCGTCTGCGGTATGTGGAAAGGTTATGGCTGTAGTTGTGATCAACTCCGCGAACCCATGCTTCAGTCAGCTGATGCACAATCGTTTTTAAACGGGTTTGCGGTGTAAGTGCAGCCCGTCTTACACCGTGCGGCACAGGCACTAGTACTGATGTCGTATACAGGGCTTTTGACATCTACAATGATAAAGTAGCTGGTTTTGCTAAATTCCTAAAAACTAATTGTTGTCGCTTCCAAGAAAAGGACGAAGATGACAATTTAATTGATTCTTACTTTGTAGTTAAGAGACACACTTTCTCTAACTACCAACATGAAGAAACAATTTATAATTTACTTAAGGATTGTCCAGCTGTTGCTAAACATGACTTCTTTAAGTTTAGAATAGACGGTGACATGGTACCACATATATCACGTCAACGTCTTACTAAATACACAATGGCAGACCTCGTCTATGCTTTAAGGCATTTTGATGAAGGTAATTGTGACACATTAAAAGAAATACTTGTCACATACAATTGTTGTGATGATGATTATTTCAATAAAAAGGACTGGTATGATTTTGTAGAAAACCCAGATATATTACGCGTATACGCCAACTTAGGTGAACGTGTACGCCAAGCTTTGTTAAAAACAGTACAATTCTGTGATGCCATGCGAAATGCTGGTATTGTTGGTGTACTGACATTAGATAATCAAGATCTCAATGGTAACTGGTATGATTTCGGTGATTTCATACAAACCACGCCAGGTAGTGGAGTTCCTGTTGTAGATTCTTATTATTCATTGTTAATGCCTATATTAACCTTGACCAGGGCTTTAACTGCAGAGTCACATGTTGACACTGACTTAACAAAGCCTTACATTAAGTGGGATTTGTTAAAATATGACTTCACGGAAGAGAGGTTAAAACTCTTTGACCGTTATTTTAAATATTGGGATCAGACATACCACCCAAATTGTGTTAACTGTTTGGATGACAGATGCATTCTGCATTGTGCAAACTTTAATGTTTTATTCTCTACAGTGTTCCCACCTACAAGTTTTGGACCACTAGTGAGAAAAATATTTGTTGATGGTGTTCCATTTGTAGTTTCAACTGGATACCACTTCAGAGAGCTAGGTGTTGTACATAATCAGGATGTAAACTTACATAGCTCTAGACTTAGTTTTAAGGAATTACTTGTGTATGCTGCTGACCCTGCTATGCACGCTGCTTCTGGTAATCTATTA**CCCGGG

Underline: *Sma*I

Bold: SARS-CoV-2, position 11664 – 14605

Fragment 7

CCCGGG**ATCAGACATACCACCCAAATTGTGTTAACTGTTTGGATGACAGATGCATTCTGCATTGTGCAAACTTTAATGTTTTATTCTCTACAGTGTTCCCACCTACAAGTTTTGGACCACTAGTGAGAAAAATATTTGTTGATGGTGTTCCATTTGTAGTTTCAACTGGATACCACTTCAGAGAGCTAGGTGTTGTACATAATCAGGATGTAAACTTACATAGCTCTAGACTTAGTTTTAAGGAATTACTTGTGTATGCTGCTGACCCTGCTATGCACGCTGCTTCTGGTAATCTATTACTAGATAAACGCACTACGTGCTTTTCAGTAGCTGCACTTACTAACAATGTTGCTTTTCAAACTGTCAAACCCGGTAATTTTAACAAAGACTTCTATGACTTTGCTGTGTCTAAGGGTTTCTTTAAGGAAGGAAGTTCTGTTGAATTAAAACACTTCTTCTTTGCTCAGGATGGTAATGCTGCTATCAGCGATTATGACTACTATCGTTATAATCTACCAACAATGTGTGATATCAGACAACTACTATTTGTAGTTGAAGTTGTTGATAAGTACTTTGATTGTTACGATGGTGGCTGTATTAATGCTAACCAAGTCATCGTCAACAACCTAGACAAATCAGCTGGTTTTCCATTTAATAAATGGGGTAAGGCTAGACTTTATTATGATTCAATGAGTTATGAGGATCAAGATGCACTTTTCGCATATACAAAACGTAATGTCATCCCTACTATAACTCAAATGAATCTTAAGTATGCCATTAGTGCAAAGAATAGAGCTCGCACCGTAGCTGGTGTCTCTATCTGTAGTACTATGACCAATAGACAGTTTCATCAAAAATTATTGAAATCAATAGCCGCCACTAGAGGAGCTACTGTAGTAATTGGAACAAGCAAATTCTATGGTGGTTGGCACAACATGTTAAAAACTGTTTATAGTGATGTAGAAAACCCTCACCTTATGGGTTGGGATTATCCTAAATGTGATAGAGCCATGCCTAACATGCTTAGAATTATGGCCTCACTTGTTCTTGCTCGCAAACATACAACGTGTTGTAGCTTGTCACACCGTTTCTATAGATTAGCTAATGAGTGTGCTCAAGTATTGAGTGAAATGGTCATGTGTGGCGGTTCACTATATGTTAAACCAGGTGGAACCTCATCAGGAGATGCCACAACTGCTTATGCTAATAGTGTTTTTAACATTTGTCAAGCTGTCACGGCCAATGTTAATGCACTTTTATCTACTGATGGTAACAAAATTGCCGATAAGTATGTCCGCAATTTACAACACAGACTTTATGAGTGTCTCTATAGAAATAGAGATGTTGACACAGACTTTGTGAATGAGTTTTACGCATATTTGCGTAAACATTTCTCAATGATGATACTCTCTGACGATGCTGTTGTGTGTTTCAATAGCACTTATGCATCTCAAGGTCTAGTGGCTAGCATAAAGAACTTTAAGTCAGTTCTTTATTATCAAAACAATGTTTTTATGTCTGAAGCAAAATGTTGGACTGAGACTGACCTTACTAAAGGACCTCATGAATTTTGCTCTCAACATACAATGCTAGTTAAACAGGGTGATGATTATGTGTACCTTCCTTACCCAGATCCATCAAGAATCCTAGGGGCCGGCTGTTTTGTAGATGATATCGTAAAAACAGATGGTACACTTATGATTGAACGGTTCGTGTCTTTAGCTATAGATGCTTACCCACTTACTAAACATCCTAATCAGGAGTATGCTGATGTCTTTCATTTGTACTTACAATACATAAGAAAGCTACATGATGAGTTAACAGGACACATGTTAGACATGTATTCTGTTATGCTTACTAATGATAACACTTCAAGGTATTGGGAACCTGAGTTTTATGAGGCTATGTACACACCGCATACAGTCTTACAGGCTGTTGGGGCTTGTGTTCTTTGCAATTCACAGACTTCATTAAGATGTGGTGCTTGCATACGTAGACCATTCTTATGTTGTAAATGCTGTTACGACCATGTCATATCAACATCACATAAATTAGTCTTGTCTGTTAATCCGTATGTTTGCAATGCTCCAGGTTGTGATGTCACAGATGTGACTCAACTTTACTTAGGAGGTATGAGCTATTATTGTAAATCACATAAACCACCCATTAGTTTTCCATTGTGTGCTAATGGACAAGTTTTTGGTTTATATAAAAATACATGTGTTGGTAGCGATAATGTTACTGACTTTAATGCAATTGCAACATGTGACTGGACAAATGCTGGTGATTACATTTTAGCTAACACCTGTACTGAAAGACTCAAGCTTTTTGCAGCAGAAACGCTCAAAGCTACTGAGGAGACATTTAAACTGTCTTATGGTATTGCTACTGTACGTGAAGTGCTGTCTGACAGAGAATTACATCTTTCATGGGAAGTTGGTAAACCTAGACCACCACTTAACCGAAATTATGTCTTTACTGGTTATCGTGTAACTAAAAACAGTAAAGTACAAATAGGAGAGTACACCTTTGAAAAAGGTGACTATGGTGATGCTGTTGTTTACCGAGGTACAACAACTTACAAATTAAATGTTGGTGATTATTTTGTGCTGACATCACATACAGTAATGCCATTAAGTGCACCTACACTAGTGCCACAAGAGCACTATGTTAGAATTACTGGCTTATACCCAACACTCAATATCTCAGATGAGTTTTCTAGCAATGTTGCAAATTATCAAAAGGTTGGTATGCAAAAGTATTCTACACTCCAGGGACCACCTGGTACTGGTAAGAGTCATTTTGCTATTGGCCTAGCTCTCTACTACCCTTCTGCTCGCATAGTGTATACAGCTTGCTCTCATGCCGCTGTTGATGCACTATGTGAGAAGGCATTAAAATATTTGCCTATAGATAAATGTAGTAGAATTATACCTGCACGTGCTCGTGTAGAGTGTTTTGATAAATTCAAAGTGAATTCAACATTAGAACAGTATGTCTTTTGTACTGTAAATGCATTGCCTGAGACGACAGCAGATATAGTTGTCTTTGATGAAATTTCAATGGCCACAAATTATGATTTGAGTGTTGTCAATGCCAGATTACGTGCTAAGCACTATGTGTACATTGGCGACCCTGCTCAATTACCTGCACCACGCACATTGCTAACTAAGGGCACACTAGAACCAGAATATTTCAATTCAGTGTGTAGACTTATGAAAACTATAGGTCCAGACATGTTCCTCGGAACTTGTCGGCGTTGTCCTGCTGAAATTGTTGACACTGTGAGTGCTTTGGTTTATGATAATAAGCTTAAAGCACATAAAGACAAATCAGCTCAATGCTTTAAAATGTTTTATAAGGGTGTTATCACGCATGATGTTTCATCTGCAA**CCCGGG

Underline: *Sma*I

Bold: SARS-CoV-2, position 14311 – 17698

Fragment 8

CCCGGG**ATGCCAGATTACGTGCTAAGCACTATGTGTACATTGGCGACCCTGCTCAATTACCTGCACCACGCACATTGCTAACTAAGGGCACACTAGAACCAGAATATTTCAATTCAGTGTGTAGACTTATGAAAACTATAGGTCCAGACATGTTCCTCGGAACTTGTCGGCGTTGTCCTGCTGAAATTGTTGACACTGTGAGTGCTTTGGTTTATGATAATAAGCTTAAAGCACATAAAGACAAATCAGCTCAATGCTTTAAAATGTTTTATAAGGGTGTTATCACGCATGATGTTTCATCTGCAATTAACAGGCCACAAATAGGCGTGGTAAGAGAATTCCTTACACGTAACCCTGCTTGGAGAAAAGCTGTCTTTATTTCACCTTATAATTCACAGAATGCTGTAGCCTCAAAGATTTTGGGACTACCAACTCAAACTGTTGATTCATCACAGGGCTCAGAATATGACTATGTCATATTCACTCAAACCACTGAAACAGCTCACTCTTGTAATGTAAACAGATTTAATGTTGCTATTACCAGAGCAAAAGTAGGCATACTTTGCATAATGTCTGATAGAGACCTTTATGACAAGTTGCAATTTACAAGTCTTGAAATTCCACGTAGGAATGTGGCAACTTTACAAGCTGAAAATGTAACAGGACTCTTTAAAGATTGTAGTAAGGTAATCACTGGGTTACATCCTACACAGGCACCTACACACCTCAGTGTTGACACTAAATTCAAAACTGAAGGTTTATGTGTTGACATACCTGGCATACCTAAGGACATGACCTATAGAAGACTCATCTCTATGATGGGTTTTAAAATGAATTATCAAGTTAATGGTTACCCTAACATGTTTATCACCCGCGAAGAAGCTATAAGACATGTACGTGCATGGATTGGCTTCGATGTCGAGGGGTGTCATGCTACTAGAGAAGCTGTTGGTACCAATTTACCTTTACAGCTAGGTTTTTCTACAGGTGTTAACCTAGTTGCTGTACCTACAGGTTATGTTGATACACCTAATAATACAGATTTTTCCAGAGTTAGTGCTAAACCACCGCCTGGAGATCAATTTAAACACCTCATACCACTTATGTACAAAGGACTTCCTTGGAATGTAGTGCGTATAAAGATTGTACAAATGTTAAGTGACACACTTAAAAATCTCTCTGACAGAGTCGTATTTGTCTTATGGGCACATGGCTTTGAGTTGACATCTATGAAGTATTTTGTGAAAATAGGACCTGAGCGCACCTGTTGTCTATGTGATAGACGTGCCACATGCTTTTCCACTGCTTCAGACACTTATGCCTGTTGGCATCATTCTATTGGATTTGATTACGTCTATAATCCGTTTATGATTGATGTTCAACAATGGGGTTTTACAGGTAACCTACAAAGCAACCATGATCTGTATTGTCAAGTCCATGGTAATGCACATGTAGCTAGTTGTGATGCAATCATGACTAGGTGTCTAGCTGTCCACGAGTGCTTTGTTAAGCGTGTTGACTGGACTATTGAATATCCTATAATTGGTGATGAACTGAAGATTAATGCGGCTTGTAGAAAGGTTCAACACATGGTTGTTAAAGCTGCATTATTAGCAGACAAATTCCCAGTTCTTCACGACATTGGTAACCCTAAAGCTATTAAGTGTGTACCTCAAGCTGATGTAGAATGGAAGTTCTATGATGCACAGCCTTGTAGTGACAAAGCTTATAAAATAGAAGAATTATTCTATTCTTATGCCACACATTCTGACAAATTCACAGATGGTGTATGCCTATTTTGGAATTGCAATGTCGATAGATATCCTGCTAATTCCATTGTTTGTAGATTTGACACTAGAGTGCTATCTAACCTTAACTTGCCTGGTTGTGATGGTGGCAGTTTGTATGTAAATAAACATGCATTCCACACACCAGCTTTTGATAAAAGTGCTTTTGTTAATTTAAAACAATTACCATTTTTCTATTACTCTGACAGTCCATGTGAGTCTCATGGAAAACAAGTAGTGTCAGATATAGATTATGTACCACTAAAGTCTGCTACGTGTATAACACGTTGCAATTTAGGTGGTGCTGTCTGTAGACATCATGCTAATGAGTACAGATTGTATCTCGATGCTTATAACATGATGATCTCAGCTGGCTTTAGCTTGTGGGTTTACAAACAATTTGATACTTATAACCTCTGGAACACTTTTACAAGACTTCAGAGTTTAGAAAATGTGGCTTTTAATGTTGTAAATAAGGGACACTTTGATGGACAACAGGGTGAAGTACCAGTTTCTATCATTAATAACACTGTTTACACAAAAGTTGATGGTGTTGATGTAGAATTGTTTGAAAATAAAACAACATTACCTGTTAATGTAGCATTTGAGCTTTGGGCTAAGCGCAACATTAAACCAGTACCAGAGGTGAAAATACTCAATAATTTGGGTGTGGACATTGCTGCTAATACTGTGATCTGGGACTACAAAAGAGATGCTCCAGCACATATATCTACTATTGGTGTTTGTTCTATGACTGACATAGCCAAGAAACCAACTGAAACGATTTGTGCACCACTCACTGTCTTTTTTGATGGTAGAGTTGATGGTCAAGTAGACTTATTTAGAAATGCCCGTAATGGTGTTCTTATTACAGAAGGTAGTGTTAAAGGTTTACAACCATCTGTAGGTCCCAAACAAGCTAGTCTTAATGGAGTCACATTAATTGGAGAAGCCGTAAAAACACAGTTCAATTATTATAAGAAAGTTGATGGTGTTGTCCAACAATTACCTGAAACTTACTTTACTCAGAGTAGAAATTTACAAGAATTTAAACCCAGGAGTCAAATGGAAATTGATTTCTTAGAATTAGCTATGGATGAATTCATTGAACGGTATAAATTAGAAGGCTATGCCTTCGAACATATCGTTTATGGAGATTTTAGTCATAGTCAGTTAGGT**CCCGGG

Underline: *Sma*I

Bold: SARS-CoV-2, position 17399 – 20358

Fragment 9

CCCG**GGAGTCACATTAATTGGAGAAGCCGTAAAAACACAGTTCAATTATTATAAGAAAGTTGATGGTGTTGTCCAACAATTACCTGAAACTTACTTTACTCAGAGTAGAAATTTACAAGAATTTAAACCCAGGAGTCAAATGGAAATTGATTTCTTAGAATTAGCTATGGATGAATTCATTGAACGGTATAAATTAGAAGGCTATGCCTTCGAACATATCGTTTATGGAGATTTTAGTCATAGTCAGTTAGGTGGTTTACATCTACTGATTGGACTAGCTAAACGTTTTAAGGAATCACCTTTTGAATTAGAAGATTTTATTCCTATGGACAGTACAGTTAAAAACTATTTCATAACAGATGCGCAAACAGGTTCATCTAAGTGTGTGTGTTCTGTTATTGATTTATTACTTGATGATTTTGTTGAAATAATAAAATCCCAAGATTTATCTGTAGTTTCTAAGGTTGTCAAAGTGACTATTGACTATACAGAAATTTCATTTATGCTTTGGTGTAAAGATGGCCATGTAGAAACATTTTACCCAAAATTACAATCTAGTCAAGCGTGGCAACCGGGTGTTGCTATGCCTAATCTTTACAAAATGCAAAGAATGCTATTAGAAAAGTGTGACCTTCAAAATTATGGTGATAGTGCAACATTACCTAAAGGCATAATGATGAATGTCGCAAAATATACTCAACTGTGTCAATATTTAAACACATTAACATTAGCTGTACCCTATAATATGAGAGTTATACATTTTGGTGCTGGTTCTGATAAAGGAGTTGCACCAGGTACAGCTGTTTTAAGACAGTGGTTGCCTACGGGTACGCTGCTTGTCGATTCAGATCTTAATGACTTTGTCTCTGATGCAGATTCAACTTTGATTGGTGATTGTGCAACTGTACATACAGCTAATAAATGGGATCTCATTATTAGTGATATGTACGACCCTAAGACTAAAAATGTTACAAAAGAAAATGACTCTAAAGAGGGTTTTTTCACTTACATTTGTGGGTTTATACAACAAAAGCTAGCTCTTGGAGGTTCCGTGGCTATAAAGATAACAGAACATTCTTGGAATGCTGATCTTTATAAGCTCATGGGACACTTCGCATGGTGGACAGCCTTTGTTACTAATGTGAATGCGTCATCATCTGAAGCATTTTTAATTGGATGTAATTATCTTGGCAAACCACGCGAACAAATAGATGGTTATGTCATGCATGCAAATTACATATTTTGGAGGAATACAAATCCAATTCAGTTGTCTTCCTATTCTTTATTTGACATGAGTAAATTTCCCCTTAAATTAAGGGGTACTGCTGTTATGTCTTTAAAAGAAGGTCAAATCAATGATATGATTTTATCTCTTCTTAGTAAAGGTAGACTTATAATTAGAGAAAACAACAGAGTTGTTATTTCTAGTGATGTTCTTGTTAACAACTAAACGAACAATGTTTGTTTTTCTTGTTTTATTGCCACTAGTCTCTAGTCAGTGTGTTAATCTTACAACCAGAACTCAATTACCCCCTGCATACACTAATTCTTTCACACGTGGTGTTTATTACCCTGACAAAGTTTTCAGATCCTCAGTTTTACATTCAACTCAGGACTTGTTCTTACCTTTCTTTTCCAATGTTACTTGGTTCCATGCTATACATGTCTCTGGGACCAATGGTACTAAGAGGTTTGATAACCCTGTCCTACCATTTAATGATGGTGTTTATTTTGCTTCCACTGAGAAGTCTAACATAATAAGAGGCTGGATTTTTGGTACTACTTTAGATTCGAAGACCCAGTCCCTACTTATTGTTAATAACGCTACTAATGTTGTTATTAAAGTCTGTGAATTTCAATTTTGTAATGATCCATTTTTGGGTGTTTATTACCACAAAAACAACAAAAGTTGGATGGAAAGTGAGTTCAGAGTTTATTCTAGTGCGAATAATTGCACTTTTGAATATGTCTCTCAGCCTTTTCTTATGGACCTTGAAGGAAAACAGGGTAATTTCAAAAATCTTAGGGAATTTGTGTTTAAGAATATTGATGGTTATTTTAAAATATATTCTAAGCACACGCCTATTAATTTAGTGCGTGATCTCCCTCAGGGTTTTTCGGCTTTAGAACCATTGGTAGATTTGCCAATAGGTATTAACATCACTAGGTTTCAAACTTTACTTGCTTTACATAGAAGTTATTTGACTCCTGGTGATTCTTCTTCAGGTTGGACAGCTGGTGCTGCAGCTTATTATGTGGGTTATCTTCAACCTAGGACTTTTCTATTAAAATATAATGAAAATGGAACCATTACAGATGCTGTAGACTGTGCACTTGACCCTCTCTCAGAAACAAAGTGTACGTTGAAATCCTTCACTGTAGAAAAAGGAATCTATCAAACTTCTAACTTTAGAGTCCAACCAACAGAATCTATTGTTAGATTTCCTAATATTACAAACTTGTGCCCTTTTGGTGAAGTTTTTAACGCCACCAGATTTGCATCTGTTTATGCTTGGAACAGGAAGAGAATCAGCAACTGTGTTGCTGATTATTCTGTCCTATATAATTCCGCATCATTTTCCACTTTTAAGTGTTATGGAGTGTCTCCTACTAAATTAAATGATCTCTGCTTTACTAATGTCTATGCAGATTCATTTGTAATTAGAGGTGATGAAGTCAGACAAATCGCTCCAGGGCAAACTGGAAAGATTGCTGATTATAATTATAAATTACCAGATGATTTTACAGGCTGCGTTATAGCTTGGAATTCTAACAATCTTGATTCTAAGGTTGGTGGTAATTATAATTACCTGTATAGATTGTTTAGGAAGTCTAATCTCAAACCTTTTGAGAGAGATATTTCAACTGAAATCTATCAGGCCGGTAGCACACCTTGTAATGGTGTTGAAGGTTTTAATTGTTACTTTCCTTTACAATCATATGGTTTCCAACCCACTAATGGTGTTGGTTACCAACCATACAGAGTAGTAGTACTTTCTTTTGAACTTCTACATGCACCAGCAACTGTTTGTGGACCTAAAAAGTCTACTAATTTGGTTAAAAACAAATGTGTCAATTTCAACTTCAATGGTTTAACAGGCACAGGTGTTCTTACTGAGTCTAACAAAAAGTTTCTGCCTTTCCAACAATTTGGCAGAGACATTGCTGACACTACTGATGCC**CGGG

Underline: *Sma*I

Bold: SARS-CoV-2, position 20110 – 23286

Fragment 10

CCCGGG**AATCTATCAGGCCGGTAGCACACCTTGTAATGGTGTTGAAGGTTTTAATTGTTACTTTCCTTTACAATCATATGGTTTCCAACCCACTAATGGTGTTGGTTACCAACCATACAGAGTAGTAGTACTTTCTTTTGAACTTCTACATGCACCAGCAACTGTTTGTGGACCTAAAAAGTCTACTAATTTGGTTAAAAACAAATGTGTCAATTTCAACTTCAATGGTTTAACAGGCACAGGTGTTCTTACTGAGTCTAACAAAAAGTTTCTGCCTTTCCAACAATTTGGCAGAGACATTGCTGACACTACTGATGCTGTCCGTGATCCACAGACACTTGAGATTCTTGACATTACACCATGTTCTTTTGGTGGTGTCAGTGTTATAACACCAGGAACAAATACTTCTAACCAGGTTGCTGTTCTTTATCAGGATGTTAACTGCACAGAAGTCCCTGTTGCTATTCATGCAGATCAACTTACTCCTACTTGGCGTGTTTATTCTACAGGTTCTAATGTTTTTCAAACACGTGCAGGCTGTTTAATAGGGGCTGAACATGTCAACAACTCATATGAGTGTGACATACCCATTGGTGCAGGTATATGCGCTAGTTATCAGACTCAGACTAATTCTCCTCGGCGGGCACGTAGTGTAGCTAGTCAATCCATCATTGCCTACACTATGTCACTTGGTGCAGAAAATTCAGTTGCTTACTCTAATAACTCTATTGCCATACCCACAAATTTTACTATTAGTGTTACCACAGAAATTCTACCAGTGTCTATGACCAAGACATCAGTAGATTGTACAATGTACATTTGTGGTGATTCAACTGAATGCAGCAATCTTTTGTTGCAATATGGCAGTTTTTGTACACAATTAAACCGTGCTTTAACTGGAATAGCTGTTGAACAAGACAAAAACACCCAAGAAGTTTTTGCACAAGTCAAACAAATTTACAAAACACCACCAATTAAAGATTTTGGTGGTTTTAATTTTTCACAAATATTACCAGATCCATCAAAACCAAGCAAGAGGTCATTTATTGAAGATCTACTTTTCAACAAAGTGACACTTGCAGATGCTGGCTTCATCAAACAATATGGTGATTGCCTTGGTGATATTGCTGCTAGAGACCTCATTTGTGCACAAAAGTTTAACGGCCTTACTGTTTTGCCACCTTTGCTCACAGATGAAATGATTGCTCAATACACTTCTGCACTGTTAGCGGGTACAATCACTTCTGGTTGGACCTTTGGTGCAGGTGCTGCATTACAAATACCATTTGCTATGCAAATGGCTTATAGGTTTAATGGTATTGGAGTTACACAGAATGTTCTCTATGAGAACCAAAAATTGATTGCCAACCAATTTAATAGTGCTATTGGCAAAATTCAAGACTCACTTTCTTCCACAGCAAGTGCACTTGGAAAACTTCAAGATGTGGTCAACCAAAATGCACAAGCTTTAAACACGCTTGTTAAACAACTTAGCTCCAATTTTGGTGCAATTTCAAGTGTTTTAAATGATATCCTTTCACGTCTTGACAAAGTTGAGGCTGAAGTGCAAATTGATAGGTTGATCACAGGCAGACTTCAAAGTTTGCAGACATATGTGACTCAACAATTAATTAGAGCTGCAGAAATCAGAGCTTCTGCTAATCTTGCTGCTACTAAAATGTCAGAGTGTGTACTTGGACAATCAAAAAGAGTTGATTTTTGTGGAAAGGGCTATCATCTTATGTCCTTCCCTCAGTCAGCACCTCATGGTGTAGTCTTCTTGCATGTGACTTATGTCCCTGCACAAGAAAAGAACTTCACAACTGCTCCTGCCATTTGTCATGATGGAAAAGCACACTTTCCTCGTGAAGGTGTCTTTGTTTCAAATGGCACACACTGGTTTGTAACACAAAGGAATTTTTATGAACCACAAATCATTACTACAGACAACACATTTGTGTCTGGTAACTGTGATGTTGTAATAGGAATTGTCAACAACACAGTTTATGATCCTTTGCAACCTGAATTAGACTCATTCAAGGAGGAGTTAGATAAATATTTTAAGAATCATACATCACCAGATGTTGATTTAGGTGACATCTCTGGCATTAATGCTTCAGTTGTAAACATTCAAAAAGAAATTGACCGCCTCAATGAGGTTGCCAAGAATTTAAATGAATCTCTCATCGATCTCCAAGAACTTGGAAAGTATGAGCAGTATATAAAATGGCCATGGTACATTTGGCTAGGTTTTATAGCTGGCTTGATTGCCATAGTAATGGTGACAATTATGCTTTGCTGTATGACCAGTTGCTGTAGTTGTCTCAAGGGCTGTTGTTCTTGTGGATCCTGCTGCAAATTTGATGAAGACGACTCTGAGCCAGTGCTCAAAGGAGTCAAATTACATTACACATAAACGAACTTATGGATTTGTTTATGAGAATCTTCACAATTGGAACTGTAACTTTGAAGCAAGGTGAAATCAAGGATGCTACTCCTTCAGATTTTGTTCGCGCTACTGCAACGATACCGATACAAGCCTCACTCCCTTTCGGATGGCTTATTGTTGGCGTTGCACTTCTTGCTGTTTTTCAGAGCGCTTCCAAAATCATAACCCTCAAAAAGAGATGGCAACTAGCACTCTCCAAGGGTGTTCACTTTGTTTGCAACTTGCTGTTGTTGTTTGTAACAGTTTACTCACACCTTTTGCTCGTTGCTGCTGGCCTTGAAGCCCCTTTTCTCTATCTTTATGCTTTAGTCTACTTCTTGCAGAGTATAAACTTTGTAAGAATAATAATGAGGCTTTGGCTTTGCTGGAAATGCCGTTCCAAAAACCCATTACTTTATGATGCCAACTATTTTCTTTGCTGGCATACTAATTGTTACGACTATTGTATACCTTACAATAGTGTAACTTCTTCAATTGTCATTACTTCAGGTGATGGCACAACAAGTCCTATTTCTGAACATGA**CCCGGG

Underline: *Sma*I

Bold: SARS-CoV-2, position 22975 – 25940

Fragment 11

CCCGG**GATGGCAACTAGCACTCTCCAAGGGTGTTCACTTTGTTTGCAACTTGCTGTTGTTGTTTGTAACAGTTTACTCACACCTTTTGCTCGTTGCTGCTGGCCTTGAAGCCCCTTTTCTCTATCTTTATGCTTTAGTCTACTTCTTGCAGAGTATAAACTTTGTAAGAATAATAATGAGGCTTTGGCTTTGCTGGAAATGCCGTTCCAAAAACCCATTACTTTATGATGCCAACTATTTTCTTTGCTGGCATACTAATTGTTACGACTATTGTATACCTTACAATAGTGTAACTTCTTCAATTGTCATTACTTCAGGTGATGGCACAACAAGTCCTATTTCTGAACATGACTACCAGATTGGTGGTTATACTGAAAAATGGGAATCTGGAGTAAAAGACTGTGTTGTATTACACAGTTACTTCACTTCAGACTATTACCAGCTGTACTCAACTCAATTGAGTACAGACACTGGTGTTGAACATGTTACCTTCTTCATCTACAATAAAATTGTTGATGAGCCTGAAGAACATGTCCAAATTCACACAATCGACGGTTCATCCGGAGTTGTTAATCCAGTAATGGAACCAATTTATGATGAACCGACGACGACTACTAGCGTGCCTTTGTAAGCACAAGCTGATGAGTACGAACTTATGTACTCATTCGTTTCGGAAGAGACAGGTACGTTAATAGTTAATAGCGTACTTCTTTTTCTTGCTTTCGTGGTATTCTTGCTAGTTACACTAGCCATCCTTACTGCGCTTCGATTGTGTGCGTACTGCTGCAATATTGTTAACGTGAGTCTTGTAAAACCTTCTTTTTACGTTTACTCTCGTGTTAAAAATCTGAATTCTTCTAGAGTTCCTGATCTTCTGGTCTAAACGAACTAAATATTATATTAGTTTTTCTGTTTGGAACTTTAATTTTAGCCATGGCAGATTCCAACGGTACTATTACCGTTGAAGAGCTTAAAAAGCTCCTTGAACAATGGAACCTAGTAATAGGTTTCCTATTCCTTACATGGATTTGTCTTCTACAATTTGCCTATGCCAACAGGAATAGGTTTTTGTATATAATTAAGTTAATTTTCCTCTGGCTGTTATGGCCAGTAACTTTAGCTTGTTTTGTGCTTGCTGCTGTTTACAGAATAAATTGGATCACCGGTGGAATTGCTATCGCAATGGCTTGTCTTGTAGGCTTGATGTGGCTCAGCTACTTCATTGCTTCTTTCAGACTGTTTGCGCGTACGCGTTCCATGTGGTCATTCAATCCAGAAACTAACATTCTTCTCAACGTGCCACTCCATGGCACTATTCTGACCAGACCGCTTCTAGAAAGTGAACTCGTAATCGGAGCTGTGATCCTTCGTGGACATCTTCGTATTGCTGGACACCATCTAGGACGCTGTGACATCAAGGACCTGCCTAAAGAAATCACTGTTGCTACATCACGAACGCTTTCTTATTACAAATTGGGAGCTTCGCAGCGTGTAGCAGGTGACTCAGGTTTTGCTGCATACAGTCGCTACAGGATTGGCAACTATAAATTAAACACAGACCATTCCAGTAGCAGTGACAATATTGCTTTGCTTGTACAGTAAGTGACAACAGATGTTTCATCTCGTTGACTTTCAGGTTACTATAGCAGAGATATTACTAATTATTATGAGGACTTTTAAAGTTTCCATTTGGAATCTTGATTACATCATAAACCTCATAATTAAAAATTTATCTAAGTCACTAACTGAGAATAAATATTCTCAATTAGATGAAGAGCAACCAATGGAGATTGATTAAACGAACATGAAAATTATTCTTTTCTTGGCACTGATAACACTCGCTACTTGTGAGCTTTATCACTACCAAGAGTGTGTTAGAGGTACAACAGTACTTTTAAAAGAACCTTGCTCTTCTGGAACATACGAGGGCAATTCACCATTTCATCCTCTAGCTGATAACAAATTTGCACTGACTTGCTTTAGCACTCAATTTGCTTTTGCTTGTCCTGACGGCGTAAAACACGTCTATCAGTTACGTGCCAGATCAGTTTCACCTAAACTGTTCATCAGACAAGAGGAAGTTCAAGAACTTTACTCTCCAATTTTTCTTATTGTTGCGGCAATAGTGTTTATAACACTTTGCTTCACACTCAAAAGAAAGACAGAATGATTGAACTTTCATTAATTGACTTCTATTTGTGCTTTTTAGCCTTTCTGCTATTCCTTGTTTTAATTATGCTTATTATCTTTTGGTTCTCACTTGAACTGCAAGATCATAATGAAACTTGTCACGCCTAAACGAACATGAAATTTCTTGTTTTCTTAGGAATCATCACAACTGTAGCTGCATTTCACCAAGAATGTAGTTTACAGTCATGTACTCAACATCAACCATATGTAGTTGATGACCCGTGTCCTATTCACTTCTATTCTAAATGGTATATTAGAGTAGGAGCTAGAAAATCAGCACCTTTAATTGAATTGTGCGTGGATGAGGCTGGTTCTAAATCACCCATTCAGTACATCGATATCGGTAATTATACAGTTTCCTGTTTACCTTTTACAATTAATTGCCAGGAACCTAAATTGGGTAGTCTTGTAGTGCGTTGTTCGTTCTATGAAGACTTTTTAGAGTATCATGACGTTCGTGTTGTTTTAGATTTCATCTAAACGAACAAACTAAAATGTCTGATAATGGACCCCAAAATCAGCGAAATGCACCCCGCATTACGTTTGGTGGACCCTCAGATTCAACTGGCAGTAACCAGAATGGAGAACGCAGTGGGGCGCGATCAAAACAACGTCGGCCCCAAGGTTTACCCAATAATACTGCGTCTTGGTTCACCGCTCTCACTCAACATGGCAAGGAAGACCTTAAATTCCCTCGAGGACAAGGCGTTCCAATTAACACCAATAGCAGTCCAGATGACCAAATTGGCTACTACCGAAGAGCTACCAGACGAATTCGTGGTGGTGACGGTAAAATGAAAGATCTCAGTCCAAGATGGTATTTCTACTACCTAGGAACTGGGCCAGAAGCTGGACTTCCCTATGGTGCTAACAAAGACGGCATCATATGGGTTGCAACTGAGGGAGCCTTGAATACACCAAAAGATCACATTGGCACCCGCAATCCTGCTAACAATGCTGCAATCGTGCTACAACTTCCTCAAGGAACAACATTGCCAAA**CCCGGG

Underline: *Sma*I

Bold: SARS-CoV-2, position 25595 – 28779

Fragment 12

CCCGGG**ATGTCTGATAATGGACCCCAAAATCAGCGAAATGCACCCCGCATTACGTTTGGTGGACCCTCAGATTCAACTGGCAGTAACCAGAATGGAGAACGCAGTGGGGCGCGATCAAAACAACGTCGGCCCCAAGGTTTACCCAATAATACTGCGTCTTGGTTCACCGCTCTCACTCAACATGGCAAGGAAGACCTTAAATTCCCTCGAGGACAAGGCGTTCCAATTAACACCAATAGCAGTCCAGATGACCAAATTGGCTACTACCGAAGAGCTACCAGACGAATTCGTGGTGGTGACGGTAAAATGAAAGATCTCAGTCCAAGATGGTATTTCTACTACCTAGGAACTGGGCCAGAAGCTGGACTTCCCTATGGTGCTAACAAAGACGGCATCATATGGGTTGCAACTGAGGGAGCCTTGAATACACCAAAAGATCACATTGGCACCCGCAATCCTGCTAACAATGCTGCAATCGTGCTACAACTTCCTCAAGGAACAACATTGCCAAAAGGCTTCTACGCAGAAGGGAGCAGAGGCGGCAGTCAAGCCTCTTCTCGTTCCTCATCACGTAGTCGCAACAGTTCAAGAAATTCAACTCCAGGCAGCAGTAGGGGAACTTCTCCTGCTAGAATGGCTGGCAATGGCGGTGATGCTGCTCTTGCTTTGCTGCTGCTTGACAGATTGAACCAGCTTGAGAGCAAAATGTCTGGTAAAGGCCAACAACAACAAGGCCAAACTGTCACTAAGAAATCTGCTGCTGAGGCTTCTAAGAAGCCTCGGCAAAAACGTACTGCCACTAAAGCATACAATGTAACACAAGCTTTCGGCAGACGTGGTCCAGAACAAACCCAAGGAAATTTTGGGGACCAGGAACTAATCAGACAAGGAACTGATTACAAACATTGGCCGCAAATTGCACAATTTGCCCCCAGCGCTTCAGCGTTCTTCGGAATGTCGCGCATTGGCATGGAAGTCACACCTTCGGGAACGTGGTTGACCTACACAGGTGCCATCAAATTGGATGACAAAGATCCAAATTTCAAAGATCAAGTCATTTTGCTGAATAAGCATATTGACGCATACAAAACATTCCCACCAACAGAGCCTAAAAAGGACAAAAAGAAGAAGGCTGATGAAACTCAAGCCTTACCGCAGAGACAGAAGAAACAGCAAACTGTGACTCTTCTTCCTGCTGCAGATTTGGATGATTTCTCCAAACAATTGCAACAATCCATGAGCAGTGCTGACTCAACTCAGGCCTAAACTCATGCAGACCACACAAGGCAGATGGGCTATATAAACGTTTTCGCTTTTCCGTTTACGATATATAGTCTACTCTTGTGCAGAATGAATTCTCGTAACTACATAGCACAAGTAGATGTAGTTAACTTTAATCTCACATAGCAATCTTTAATCAGTGTGTAACATTAGGGAGGACTTGAAAGAGCCACCACATTTTCACCGAGGCCACGCGGAGTACGATCGAGTGTACAGTGAACAATGCTAGGGAGAGCTGCCTATATGGAAGAGCCCTAATGTGTAAAATTAATTTTAGTAGTGCTATCCCCATGTGATTTTAATAGCTTCTTAGGAGAATGAC**aaaaaaaaaaaaaaaaaaaaaaaaaaaaa***CGGCCG*ggccggcatggtcccagcctcctcgctggcgccggctgggcaacattccgaggggaccgtcccctcggtaatggcgaatgggacgggccctgcgatatcgcgacgaggatctagatcctctagagtcgacctgcaggcatgcaagcttgagtattctatagtctcacctaaatagcttgg**CCCGGG

Underline: *Sma*I

Bold: SARS-CoV-2, position 28274 – 29870

Lowercase: poly(A) tail

Underline, bold, italic: *Eag*I

Lowercase, bold: Hepatitis D virus ribozyme and pCC1 sequences

Modification of SARS-CoV-2 synthetic fragment 11 to introduce TurboGFP transgene

Fragment 13

***GATGGCAACTAGCACTCTCC*AAGGGTGTTCACTTTGTTTGCAACTTGCTGTTGTTGTTTGTAACAGTTTACTCACACCTTTTGCTCGTTGCTGCTGGCCTTGAAGCCCCTTTTCTCTATCTTTATGCTTTAGTCTACTTCTTGCAGAGTATAAACTTTGTAAGAATAATAATGAGGCTTTGGCTTTGCTGGAAATGCCGTTCCAAAAACCCATTACTTTATGATGCCAACTATTTTCTTTGCTGGCATACTAATTGTTACGACTATTGTATACCTTACAATAGTGTAACTTCTTCAATTGTCATTACTTCAGGTGATGGCACAACAAGTCCTATTTCTGAACATGACTACCAGATTGGTGGTTATACTGAAAAATGGGAATCTGGAGTAAAAGACTGTGTTGTATTACACAGTTACTTCACTTCAGACTATTACCAGCTGTACTCAACTCAATTGAGTACAGACACTGGTGTTGAACATGTTACCTTCTTCATCTACAATAAAATTGTTGATGAGCCTGAAGAACATGTCCAAATTCACACAATCGACGGTTCATCCGGAGTTGTTAATCCAGTAATGGAACCAATTTATGATGAACCGACGACGACTACTAGCGTGCCTTTGTAAGCACAAGCTGATGAGTACGAACTTATGTACTCATTCGTTTCGGAAGAGACAGGTACGTTAATAGTTAATAGCGTACTTCTTTTTCTTGCTTTCGTGGTATTCTTGCTAGTTACACTAGCCATCCTTACTGCGCTTCGATTGTGTGCGTACTGCTGCAATATTGTTAACGTGAGTCTTGTAAAACCTTCTTTTTACGTTTACTCTCGTGTTAAAAATCTGAATTCTTCTAGAGTTCCTGATCTTCTGGTCTAAACGAACTAAATATTATATTAGTTTTTCTGTTTGGAACTTTAATTTTAGCCATGGCAGATTCCAACGGTACTATTACCGTTGAAGAGCTTAAAAAGCTCCTTGAACAATGGAACCTAGTAATAGGTTTCCTATTCCTTACATGGATTTGTCTTCTACAATTTGCCTATGCCAACAGGAATAGGTTTTTGTATATAATTAAGTTAATTTTCCTCTGGCTGTTATGGCCAGTAACTTTAGCTTGTTTTGTGCTTGCTGCTGTTTACAGAATAAATTGGATCACCGGTGGAATTGCTATCGCAATGGCTTGTCTTGTAGGCTTGATGTGGCTCAGCTACTTCATTGCTTCTTTCAGACTGTTTGCGCGTACGCGTTCCATGTGGTCATTCAATCCAGAAACTAACATTCTTCTCAACGTGCCACTCCATGGCACTATTCTGACCAGACCGCTTCTAGAAAGTGAACTCGTAATCGGAGCTGTGATCCTTCGTGGACATCTTCGTATTGCTGGACACCATCTAGGACGCTGTGACATCAAGGACCTGCCTAAAGAAATCACTGTTGCTACATCACGAACGCTTTCTTATTACAAATTGGGAGCTTCGCAGCGTGTAGCAGGTGACTCAGGTTTTGCTGCATACAGTCGCTACAGGATTGGCAACTATAAATTAAACACAGACCATTCCAGTAGCAGTGACAATATTGCTTTGCTTGTACAGTAAGTGACAACAGATGTTTCATCTCGTTGACTTTCAGGTTACTATAGCAGAGATATTACTAATTATTATGAGGACTTTTAAAGTTTCCATTTGGAATCTTGATTACATCATAAACCTCATAATTAAAAATTTATCTAAGTCACTAACTGAGAATAAATATTCTCAATTAGATGAAGAGCAACCAATGGAGATTGATTAAACGAACATGAAAATTATTCTTTT*CTTGGCACTGATAACACTCGCT***gagagcgacgagagcggc

Underline, italic: Primer binding site

Lowercase: TurboGFP sequence, added as overhang to generate overlaps with fragment 14

Bold, with/without underline and italic: SARS-CoV-2, position 25595 - 27432

Fragment 14 containing TurboGFP gene

ttattcttttcttggcactgataacactcgct***GAGAGCGACGAGAGCGGC*CTGCCCGCCATGGAGATCGAGTGCCGCATCACCGGCACCCTGAACGGCGTGGAGTTCGAGCTGGTGGGCGGCGGAGAGGGCACCCCCGAGCAGGGCCGCATGACCAACAAGATGAAGAGCACCAAAGGCGCCCTGACCTTCAGCCCCTACCTGCTGAGCCACGTGATGGGCTACGGCTTCTACCACTTCGGCACCTACCCCAGCGGCTACGAGAACCCCTTCCTGCACGCCATCAACAACGGCGGCTACACCAACACCCGCATCGAGAAGTACGAGGACGGCGGCGTGCTGCACGTGAGCTTCAGCTACCGCTACGAGGCCGGCCGCGTGATCGGCGACTTCAAGGTGATGGGCACCGGCTTCCCCGAGGACAGCGTGATCTTCACCGACAAGATCATCCGCAGCAACGCCACCGTGGAGCACCTGCACCCCATGGGCGATAACGATCTGGATGGCAGCTTCACCCGCACCTTCAGCCTGCGCGACGGCGGCTACTACAGCTCCGTGGTGGACAGCCACATGCACTTCAAGAGCGCCATCCACCCCAGCATCCTGCAGAACGGGGGCCCCATGTTCGCCTTCCGCCGCGTGGAGGAGGATCACAGCAACACCGAGCTGGGCATCGTGGAGTACCAGCACGCCTTCAAGACCCCG*GATGCAGATGCCGGTGAAGAA*T**aactttactctccaatttttcttattgttgc

Underline italic: Primer binding site

Lowercase at the 5' end: SARS-CoV-2, position 28108 – 28130 added as overhang to generate overlaps with fragment 13

Lowercase at the 3' end: SARS-CoV-2, position 28108 – 28130 added as overhang to generate overlaps with fragment 15

Bold, with/without underline and italic: TurboGFP

Fragment 15

gcagatgccggtgaagaata**act*TTACTCTCCAATTTTTCTTATTGTTGC*GGCAATAGTGTTTATAACACTTTGCTTCACACTCAAAAGAAAGACAGAATGATTGAACTTTCATTAATTGACTTCTATTTGTGCTTTTTAGCCTTTCTGCTATTCCTTGTTTTAATTATGCTTATTATCTTTTGGTTCTCACTTGAACTGCAAGATCATAATGAAACTTGTCACGCCTAAACGAACATGAAATTTCTTGTTTTCTTAGGAATCATCACAACTGTAGCTGCATTTCACCAAGAATGTAGTTTACAGTCATGTACTCAACATCAACCATATGTAGTTGATGACCCGTGTCCTATTCACTTCTATTCTAAATGGTATATTAGAGTAGGAGCTAGAAAATCAGCACCTTTAATTGAATTGTGCGTGGATGAGGCTGGTTCTAAATCACCCATTCAGTACATCGATATCGGTAATTATACAGTTTCCTGTTTACCTTTTACAATTAATTGCCAGGAACCTAAATTGGGTAGTCTTGTAGTGCGTTGTTCGTTCTATGAAGACTTTTTAGAGTATCATGACGTTCGTGTTGTTTTAGATTTCATCTAAACGAACAAACTAAAATGTCTGATAATGGACCCCAAAATCAGCGAAATGCACCCCGCATTACGTTTGGTGGACCCTCAGATTCAACTGGCAGTAACCAGAATGGAGAACGCAGTGGGGCGCGATCAAAACAACGTCGGCCCCAAGGTTTACCCAATAATACTGCGTCTTGGTTCACCGCTCTCACTCAACATGGCAAGGAAGACCTTAAATTCCCTCGAGGACAAGGCGTTCCAATTAACACCAATAGCAGTCCAGATGACCAAATTGGCTACTACCGAAGAGCTACCAGACGAATTCGTGGTGGTGACGGTAAAATGAAAGATCTCAGTCCAAGATGGTATTTCTACTACCTAGGAACTGGGCCAGAAGCTGGACTTCCCTATGGTGCTAACAAAGACGGCATCATATGGGTTGCAACTGAGGGAGCCTTGAATACACCAAAAGATCACATTGGCACCCGCAATCCTGCTAACAATGCTGCAATCGTGCTACAACTT*CCTCAAGGAACAACATTGCCAAA***

Underline italic: Primer binding site

Lowercase: TurboGFP, added as overhang to generate overlaps with fragment 14

Bold, with/without underline and italic: SARS-CoV-2, position 28128 – 29229
